# Evolution of developmental bias explains divergent patterns of phenotypic evolution

**DOI:** 10.1101/2025.02.04.636455

**Authors:** Joao Picao-Osorio, Charlotte Bouleau, Pablo M. Gonzalez de la Rosa, Lewis Stevens, Nina Fekonja, Mark Blaxter, Christian Braendle, Marie-Anne Félix

## Abstract

Rates of phenotypic evolution vary across traits, and these evolutionary patterns themselves evolve. Understanding how development contributes—or biases—such patterns remains a challenge because it requires large-scale measurement of phenotypic variation resulting from random mutations across multiple species. Using the experimentally tractable system of nematode vulval development, we quantified the mutational sensitivity of six cell fates across two nematode clades. The results show that within each clade, mutational sensitivity is sufficient to explain the observed phenotypic evolution. The difference between clades can be explained by a simple spatial shift across cells of the sensitive region of a Wnt signaling dose-response curve. These findings underscore the importance of integrating the evolving genotype-phenotype map into the understanding of phenotypic change at both micro- and macroevolutionary scales.

## Main Text

Phenotypic traits evolve at different rates, but how different mechanisms—such as natural selection, genetic drift, mutation and development— explain these rates remain under active debate (*1*). As noted by the palaeontologist G.G. Simpson, a prominent figure of the mid-20th century evolutionary synthesis, patterns of phenotypic evolution themselves evolve, so that different clades of organisms may display distinct directions of evolutionary change (*2*). Natural selection is a central force in the evolution of phenotypes yet it only operates on available phenotypic variation. Simpson remarked, that in the space of phenotypes, “It is not only improbable but also inconceivable that mutations in every imaginable direction occur with equal frequency,” and called “to determine whether or not mutations (in the broadest sense) are directional and parallel to the actual direction of evolution” (*2*). Simpson’s question translates to quantifying the spectrum of phenotypes produced by random mutation of the genotype: in the framework of quantitative genetics, the mutational variance (*V_M_*) (*3*). Given that development mediates the relationship between genotype and phenotype (*4–6*), the non-isotropic pattern of phenotypic variation generated by random mutation is called developmental bias (*7–9*). Determining its importance in phenotypic evolution has long been recognized as the central focus of evolutionary developmental biology (*4–6, 9–13*). Yet, its challenging experimental measure has only been attempted in few systems, including dipteran wing shape (*14*), mitotic spindle traits (*15*) and *Caenorhabditis* vulva precursor cells (*16*). Furthermore, developmental bias itself may evolve, which is the focus of our study. Specifically, as Simpson predicted, developmental bias may shift over macroevolutionary timescales. Ideal systems for straightforward comparisons of different traits in distant species are those with serial homologs, such as mammalian teeth (*2, 17, 18*) or butterfly eyespots (*19–21*). However, these two systems are not well-suited for systematically assessing the effects of random mutation across distinct clades.

Here, we use the homologous cellular framework of nematode vulval development to test whether evolutionary change in developmental biases, as measured by the effect of random mutation, can explain the differences in phenotypic evolution between two clades. In a wide range of nematode species, 12 ventral epidermal precursor cells are positioned along the anterior-posterior axis of early larvae (Fig. 1A). Of those, the six cells numbered P3.p to P8.p form the vulva competence group in *Caenorhabditis elegans* (*22*). These cells undergo a series of fate specifications and divisions under the influence of several signalling pathways. The central cells P5.p, P6.p, P7.p normally form the vulva proper, required for oviposition and copulation with males (Fig. 1B, fig. S1), but can be replaced by P3.p, P4.p and P8.p. These external non-vulval cells usually divide once at the third larval stage and fuse to the epidermal syncytium. Cell fates of P4.p to P8.p are robust to environmental and genetic perturbations (*16, 23, 24*), and remain largely unchanged across the *Caenorhabditis* genus (*25–27*). In contrast, in *C. elegans* P3.p has a stochastically variable cell fate: P3.p can either divide once like P4.p or fuse directly with the syncytium during the L2 stage without division (*23, 28*). The frequency of this binary cell fate decision (L2 stage fusion versus division) evolves rapidly within and among *Caenorhabditis* species, and is highly sensitive to environmental and genetic perturbations (*16, 24, 25, 27*). In other genera of the family Rhabditidae, such as *Oscheius* (*26*), the same six cells remain unfused during early larval development (*29*) and P5.p, P6.p and P7.p progeny also form the vulva. However, in *Oscheius*, P3.p almost never divides and loses competence (*30*). In contrast, P4.p and P8.p in most species usually divide twice but can adopt one of three distinct cell division patterns (no, one, or two cell divisions), with the frequency of these patterns varying significantly both within and among *Oscheius* species and in response to environmental variation (*25*). These observations support a compelling and testable hypothesis: divergent patterns of phenotypic evolution arise as a consequence of the evolution of developmental biases.

**Figure 1.**
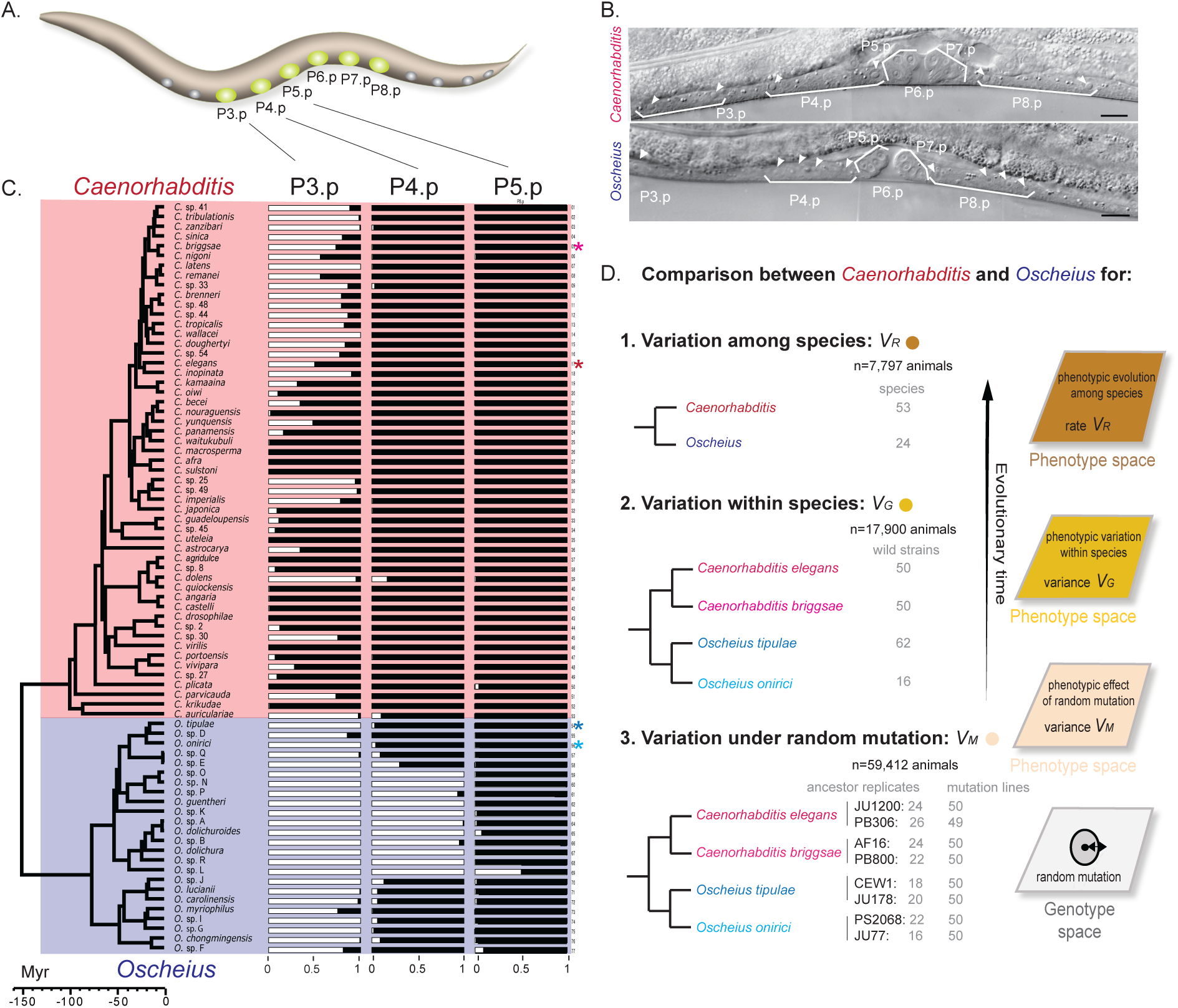
Divergent patterns of vulva fates in two nematode genera. (A) Diagram of a L1 nematode larva with the six vulva precursor cells (VPC) highlighted in green, modified from Openworm.org. Additional Pn.p cells are shown in grey. Anterior is to the left and ventral side down. (B) Nomarski images of a mid-L4 stage vulva displaying the descendants of VPCs in *C. elegans* and *O. tipulae*. In the *C. elegans* individual (top), P3.p, P4.p and P8.p all undergo a division as shown by the presence of two nuclei per mother Pn.p cell (arrowheads). In the *O. tipulae* individual (bottom), P3.p does not divide and P4.p and P8.p divide twice, producing four nuclei (arrowheads). Pictures from (*53*). Scale bar represents 10 μm. Anterior is to the left and the ventral side down. (C) Horizontal bar plots showing the fate of the P3.p, P4.p and P5.p cells plotted along the chronogram tree of *Caenorhabditis* (red) and *Oscheius* (blue). The proportion of animals with the most common type of division of P3.p, P4.p and P5.p in each genus is displayed in black. See fig. S5 and Data S1 for detailed data with all observed cell division patterns, showing further variation in *Oscheius*. Myr: Million year (estimate). Stars represent the species that were used in further studies. (D) Schematic representation of the experimental design scoring vulval cell fates in two nematode genera across three evolutionary time scales: 1. among species (*V_R_*; top); 2. within species (*V_G_*; middle); and 3. upon *de novo* mutation (*V_M_*; bottom). Numbers of lines and individuals phenotyped in the present work are indicated. Schematic genotype-phenotype maps are represented on the right.

Here we test this hypothesis using the nematode vulva precursor cell fates, in a quantative genetic framework. The overall design is shown in Fig. 1D. We represent results on the scale of phenotypes in Fig. 2 and of estimated variance in Fig. 3. For readers unfamiliar with the quantitative genetic framework, consider these variances as measures of phenotypic variation for each trait (the fate of each cell) due to genetic variation at three evolutionary scales: in response to random mutation (*M*), and in intraspecific (*G*) and interspecific (*R*) variation (Fig. 1D). Altogether, we report here on the phenotype of over 80,000 individual animals.

**Figure 2.**
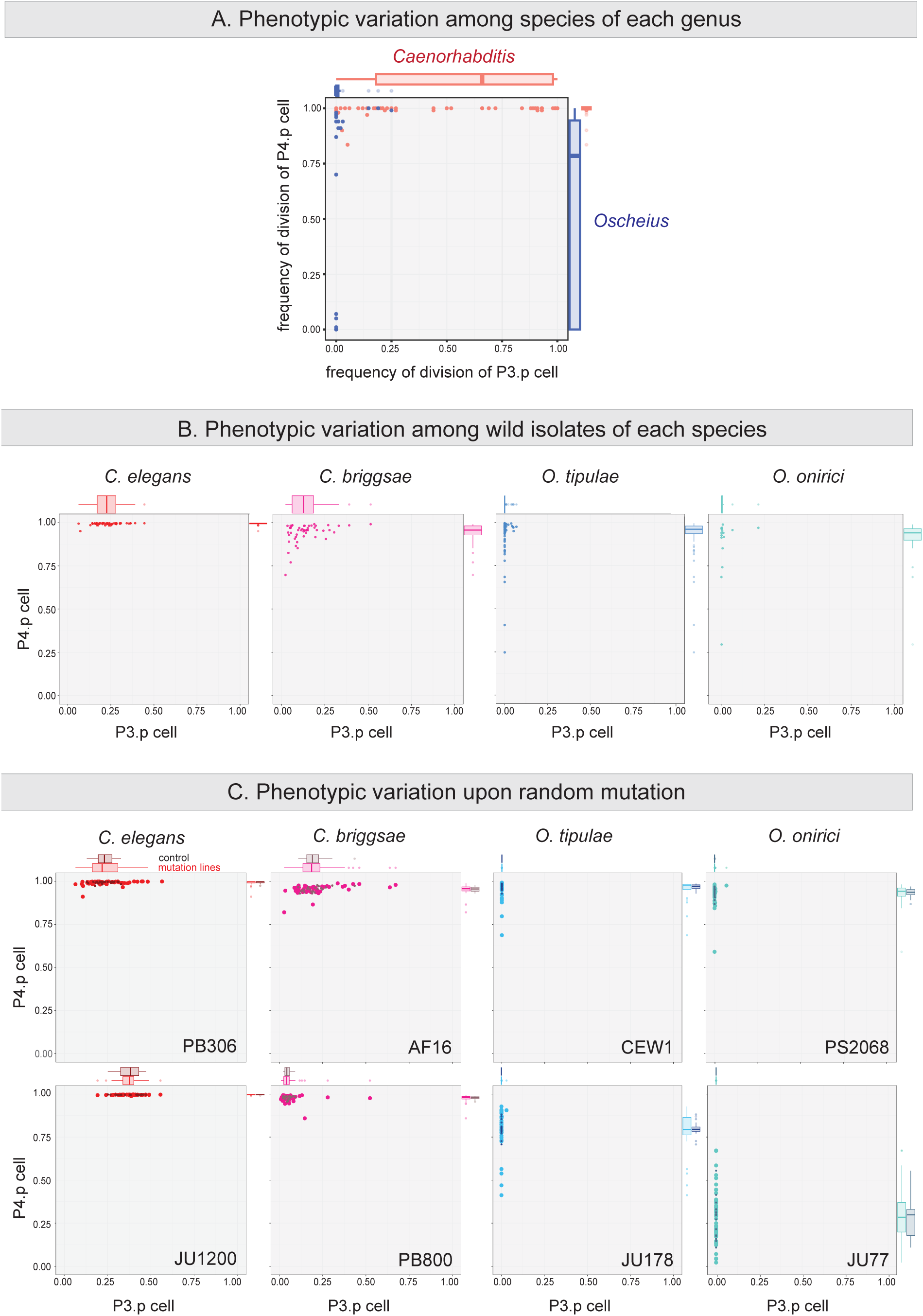
Each nematode genus shows a specific pattern of phenotypic variation for P3.p and P4.p cell fates consistent across evolutionary scales. (A) Division frequency of P3.p and P4.p in species of *Caenorhabditis* (salmon) and *Oscheius* (dark blue). Each dot denotes the average division frequency of a species. On top and on the right side of each plot are boxplots displaying the data distribution for P3.p and P4.p, respectively. (B) Division frequency of P3.p and P4.p in wild isolates of the four species as in (A). Each dot represents the Best linear unbiased prediction (BLUP) for each wild isolate. (C) Division frequency of P3.p (x-axis) and P4.p (y-axis) upon random mutation in two isolates of *C. elegans* (red), *C. briggsae* (pink), *O. tipulae* (blue) and *O. onirici* (turquoise). Each dot represents the Best linear unbiased prediction (BLUP) in the observed data scale for each strain, corrected for block and observer structure. Darker colours are the pseudo-ancestral control lines and lighter colours the random mutant lines. Numbers of phenotyped individuals and lines are provided on Fig. 1D and methods. As in Fig. 1C, is displayed the frequency of the most common type of cell lineage in each genus.

**Figure 3.**
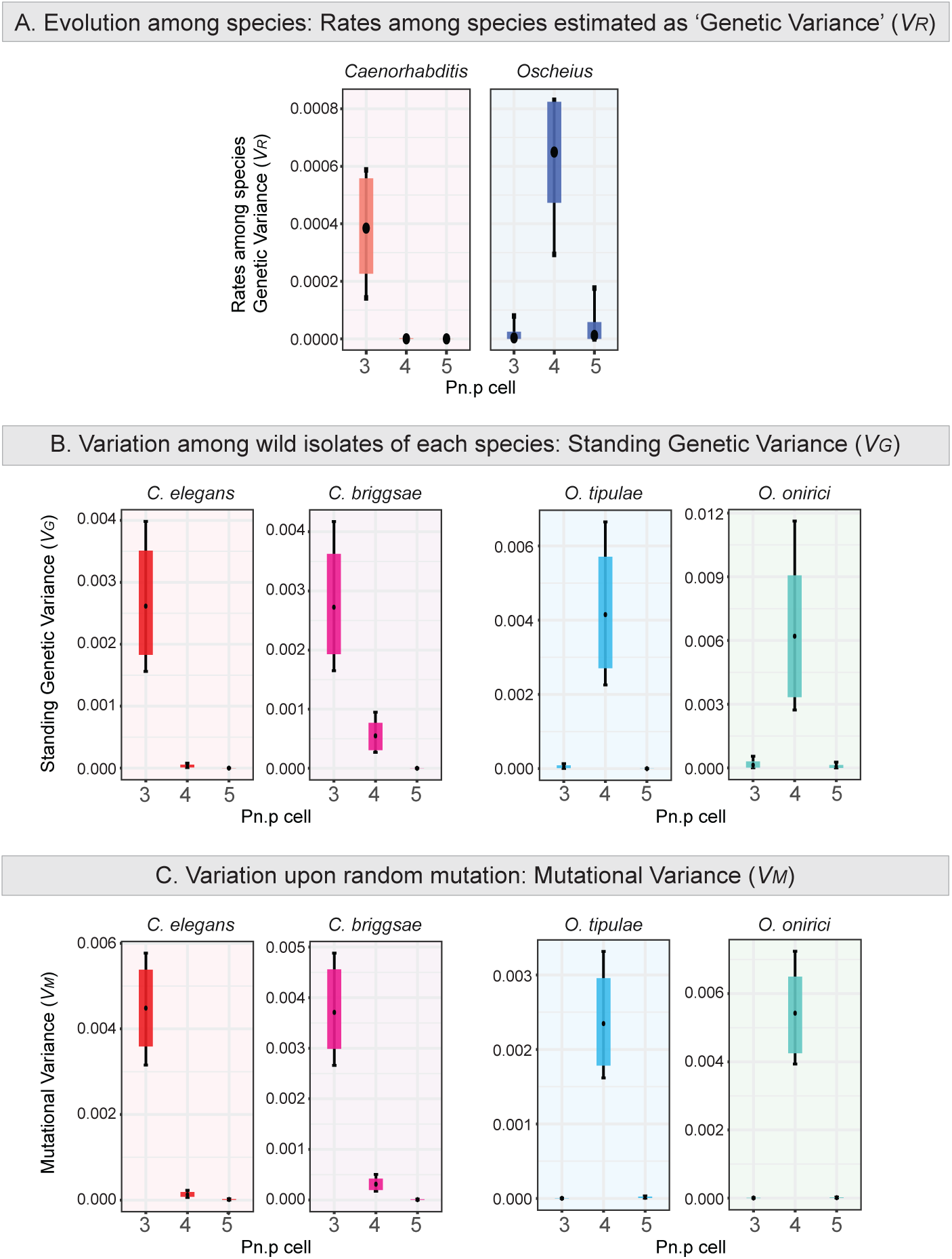
Each nematode genus shows consistent genetic variances for vulval traits across evolutionary scales. (A) Rate among species genetic variance (*V_R_*) of the three anterior VPCs (P3.p to P5.p) estimated for the *Caenorhabditis* and *Oscheius* genera. *V_R_* quantifies phenotypic evolutionary rates by estimating the increase of genetic variance among species per million years assuming a Brownian motion model of trait evolution. The scale of the genetic variance (y-axis) is the same for both graphs. (B) Standing genetic variances (*V_G_*) of the three anterior VPCs in *C. elegans* (red), *C. briggsae* (pink), *O. tipulae* (blue) and *O. onirici* (turquoise) estimated from the variation among wild isolates adjusted for the intrastrain variation of two APL controls on each species. (C) Mutational variances (*V_M_*) of the three anterior VPCs in *C. elegans* (red), *C. briggsae* (pink), *O. tipulae* (blue) and *O. onirici* (turquoise). The mutational variance is estimated as the genetic variance among random mutation lines of two isolates corrected for their ancestral pseudo-line (APL) controls. The scale of the genetic variance (y-axis) is customised to each species in (B) and (C). Each estimation displays the median (black circle), 83% (coloured bar) and 95% (black bar) credible intervals of the posterior distributions of the genetic variances in the observed data scale. Each posterior distribution has a minimum effective sample size of 2000. See fig. S17 for the estimates for the six cells.

### Divergent rates of cell fate evolution across clades

We started by determining the fates of the six vulval precursor cells using Nomarski optics in 53 *Caenorhabditis* and 24 *Oscheius* species (Fig. 1). To infer the phylogenetic relationships of these species (figs. S3-S4), we used 560 orthologues obtained from assembled genomes and transcriptomes. We calibrated the phylogenetic tree to estimate the divergence time between the *Caenorhabditis* and *Oscheius* genera at approximately 160 million years ago (MYA; Figs. 1C, S5). We then mapped the vulval cell fate changes onto this phylogeny (figs. S6). As expected, the division frequency of P3.p showed the greatest variation in *Caenorhabditis*, whereas in *Oscheius*, P4.p exhibited the most variation, producing between one and four cells (Figs. 1C, 2A, figs. S5, S6); for simplicity, we consider here only the most common pattern in *Oscheius*, where two divisions produce four cells, versus all alternative patterns. We used phylogenetic mixed models (PMM) (*31*) to quantify rates of phenotypic evolution by estimating the ‘rate among species genetic variance per million year’ (*V_R_*). In *Caenorhabditis*, the fate of P3.p evolved approximately a thousand times faster than the fate of P4.p (Fig. 3A, 4A right panel). Conversely, in *Oscheius*, the fate of P4.p evolved about a hundred times faster than that of P3.p (Fig. 3A, 4A right panel). The phylogenetic relationships within each clade explained a large proportion of the phenotypic variance of the fastest-evolving cells (fig. S8), as measured by phylogenetic heritability (*32*). Thus, the fates of P3.p and P4.p cells display contrasting patterns of phenotypic evolution between *Caenorhabditis* and *Oscheius* species.

### Alignment of intraspecific variation with evolution in each genus

Next, we asked whether we could detect corresponding patterns within species. For this, we scored in parallel wild isolates of *C. elegans*, *C. briggsae*, *O. tipulae* and *O. onirici* to estimate the genetic (*V_G_*) variances of the six traits. These four species were chosen because they reproduce primarily by selfing and each strain is composed of isogenic individuals. Phenotyping followed a block design and blinded approach with four observers (see Methods).

Similar to results for interspecific evolution, P3.p within each *Caenorhabditis* species exhibited much greater phenotypic variation in division frequency compared to any other cell (Fig. 2B; fig. S9). Specifically, *C. briggsae* and *C. elegans* displayed approximately ten and one hundred times higher *V_G_* for P3.p than for P4.p, respectively (Figs. 3B, 4A middle panel, figs. S10A, S11A). In contrast, we found that within the two *Oscheius* species, P4.p showed the greatest phenotypic variation (Fig. 2B; fig. S9). In *O. tipulae* and *O. onirici*, *V_G_* of P4.p was a hundred times greater than that of P3.p (Figs. 3B, 4A, figs. S10B, S11B). Although three of the four species hardly show any variation for one of the phenotypic axes, *C. briggsae* exhibits a more complex pattern, displaying variation along both axes, yet with a stronger effect for P3.p fate. In conclusion, the rate of cell fate evolution differs significantly between the two genera and the axis of variation within species aligns with that observed among species of the genus.

**Figure 4.**
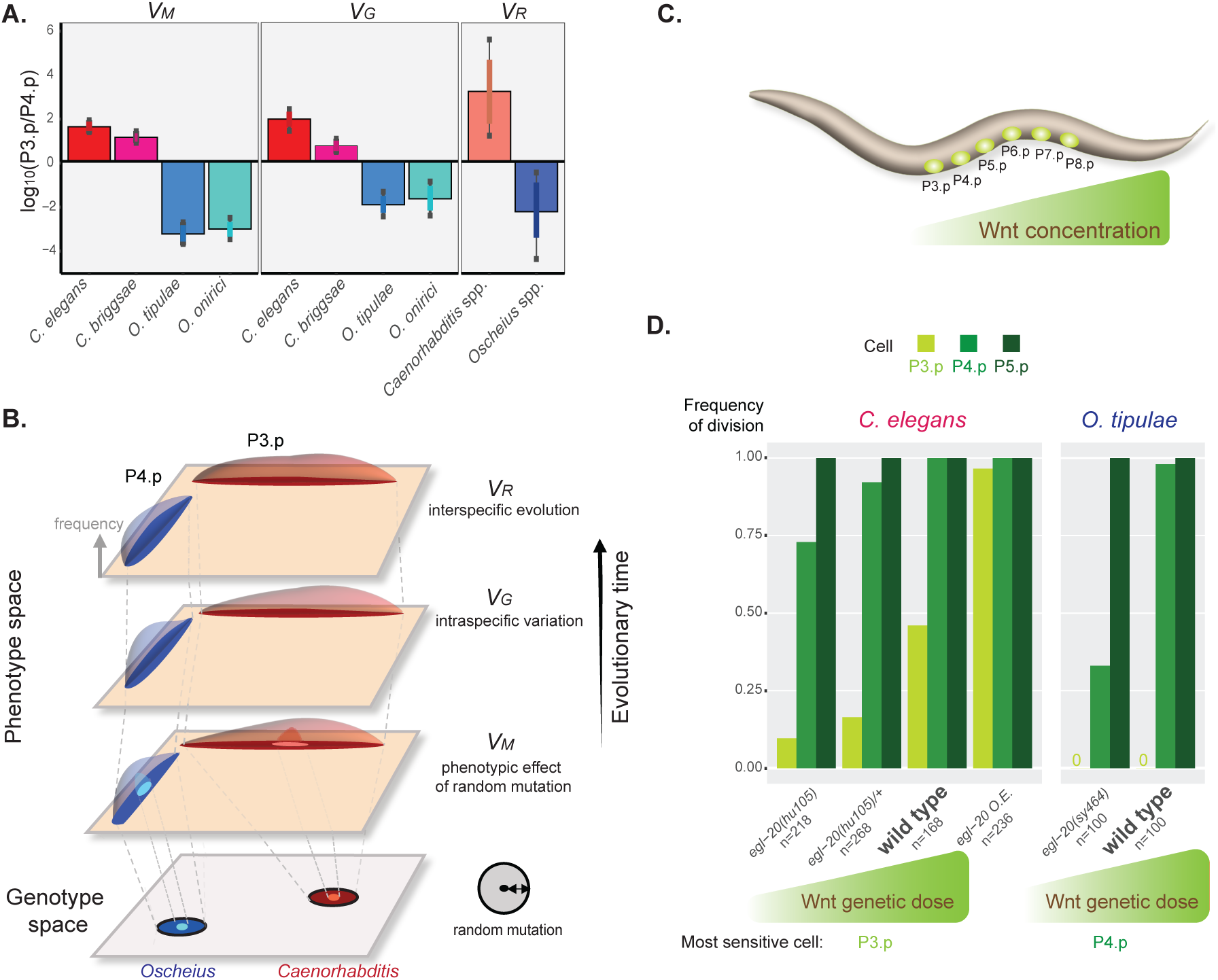
Divergence in developmental bias matches divergence in phenotypic evolutionary rates between the two nematode clades. (A) Ten-fold change of the ratio between P3.p and P4.p genetic variances across *V_M_*, *V_G_* and *V_R_* in the two genera. Bar plots display the median of the ratio of the posterior distributions of P3.p over P4.p variances, with the 83% (coloured bar) and 95% (black bar) credible intervals, plotted on a log scale. (B) Model of the genotype-phenotype map across evolutionary time scales that schematizes the data. From bottom to top, the contrasting developmental bias in *Caenorhabditis* and *Oscheius* revealed by the orthogonal phenotypic variation upon *de novo* mutation (*V_M_*) is aligned with the axes of heritable phenotypic variation within (*V_G_*) and among (*V_R_*) species of the respective genus. (C) Diagram of an early *C. elegans* larva with the six VPCs (green) and the posterior Wnt protein gradient (based on (*45*), with a simplified shape). P3.p is placed at the fading end of the gradient. (D) Bar plots showing the dose-response effects of *egl-20/Wnt* genetic modulation on the division frequency of P3.p, P4.p and P5.p in *C. elegans* (N2 genetic background) and *O. tipulae* (CEW1 genetic background). The *C. elegans* data are replotted from (*43*). For simplicity, we here neglected the few cases of miscentering of the 1° fate on P5.p. O.E.: overexpression.

### Evolutionary change in developmental bias

We next asked whether the contrasting phenotypic evolutionary rates of the six precursor cells could be explained by the evolution of developmental bias between *Caenorhabditis* and *Oscheius*. To this end, we created eight panels of lines with *de novo* mutations induced by chemical mutagenesis in wild isolates of both genera, allowing us to quantify the mutational variance (*V_M_*) of cell fates (fig. S2A). To account for genotype-specific effects on mutational variances, we performed the chemical mutagenesis on two wild isolates of two species per clade (Fig. 1D bottom panel). To account for environmental and other non-genetic effects, we phenotyped the ancestral isogenic wild isolates multiple times in parallel with the mutant lines (table S5, Data S6). The phenotyping was also performed in parallel with the wild isolates scored to estimate the standing genetic variances (*V_G_*) mentioned above.

We found that, in each genus, vulval phenotypic variation generated by *de novo* mutation is anisotropic in phenotype space (Fig. 2C, fig. S12), thus revealing a developmental bias. Moreover, this developmental bias differs between the two genera. In both *Caenorhabditis* species, the principal axis of phenotypic variation resulting from random mutation concerns P3.p fate: *C. briggsae* and *C. elegans* showed over ten times greater variation upon random mutation (*V_M_*) for P3.p compared to P4.p (Figs. 3C, 4A left panel), in line with our previous study using spontaneous mutation accumulation lines (*16*). The difference between the two cells is more nuanced in *C. briggsae* than in *C. elegans*, indicating evolution within the genus. In contrast, the two *Oscheius* species showed significantly greater phenotypic variation in P4.p fate in response to new mutations (Fig. 2C, fig. S12). P4.p presented a *V_M_* that was a thousand times greater than that of P3.p in both *O. tipulae* and *O. onirici* (Fig. 3C, 4A left panel).

The variances in fates of the two cells at the three levels (mutation, polymorphism, divergence) can be expressed as a ratio, which is robust to differences in mutation efficacy, species genetic diversity and genus age (Fig. 4A). These results demonstrate evolutionary change in developmental bias between the *Caenorhabditis* and *Oscheius* clades. Altogether, our results reveal a strong alignment between the axes of heritable phenotypic variation resulting from *de novo* mutations and the axes of natural variation within each species and genus, though they contrast between genera. Thus, the change in developmental bias predicts the axis of phenotypic evolution within each genus (Fig. 4B).

## Discussion and Conclusions

The mechanisms underlying the tempo and mode of evolution have been a subject of active debate for a long time, aiming to bridge micro- and macro-evolution (*2, 33–36*). Our results imply that the evolutionary change in mutational variances between two nematode clades can predict patterns of phenotypic evolution of the vulva precursor cell fates. This has important implications for the role of development in shaping the tempo and mode of phenotypic evolution. Over short timescales within a lineage, species with a similar developmental bias are likely to exhibit shared patterns of phenotypic evolution (*15, 37–39*). On longer timescales, lineages with distinct developmental biases may undergo divergent patterns of phenotypic evolution for different traits.

Our study cannot address evolutionary mechanisms (selection or drift) underlying the macro-evolutionary change in bias. Variation in P3.p and P4.p fates may have little impact on organismal biology because these cells (or their progeny) will fuse to the epidermal syncytium regardless of whether they divide. Their division may be selectively maintained by these cells serving as a backup system in environments that increase deviant fates for the central vulval cells (*24*): for these cells to be competent they must remain unfused at the second larval stage. However, selection pressure for the maintenance of a large competence group appears weak, since some nematodes restrict competence to only those cells that actually form the vulva (*26, 40*). Hence, the observed evolution of mutational and standing genetic variances in these cell fate traits likely results from selection on other traits, particularly those involving the same signalling pathways, such as the highly pleiotropic Wnt pathway (*41*). We consider this scenario of apparently low selection on peripheral cells of the competence group to increase the potential for developmental bias to manifest.

Mechanistically, we propose that the evolution of developmental bias in nematode vulva cell fate patterning is due to distinct developmental sensitivities of the cells, based on their anterior-posterior positioning in a Wnt gradient (Fig. 4C). Past research shows that in *C. elegans*, altering the doses of either of two genes encoding Wnt ligands, *egl-20* and *cwn-1*, significantly affects the P3.p division frequency more than that of P4.p (*42, 43*) (Fig. 4C, D). Specifically, reducing the Wnt dose leads to a lower P3.p division frequency. The two Wnt genes, especially *egl-20*, are transcribed in a posterior body region, forming a protein gradient that extends toward the anterior (*44, 45*), placing P3.p at the fading end of this gradient. Indeed, interindividual variability in downstream Wnt signalling through β-catenin/BAR-1 can be detected in P3.p (but not in P4.p), and is linked to its stochastic cell fate outcome (*46*). These observations strongly suggest that the anterior position of P3.p contributes to its heightened developmental sensitivity to stochastic, environmental, and genetic variation (*24*). In line with this, we previously showed that the high mutational variance of P3.p in *Caenorhabditis* arises from a broad mutational target, where alterations in various biochemical pathways affect its cell fate (*47*). *Oscheius tipulae*, displays an overall conserved pattern of Wnt expression (*48*), but modulation of *egl-20/Wnt* dose influences the division frequency of P4.p but not that of P3.p, which already shows no division in the wild-type reference (Fig. 4D). These results suggest that the evolution of developmental bias corresponds to the evolution of developmental sensitivity of the cells, which itself may be due to quantitative variation in the Wnt gradient or response.

This mode of phenotypic evolution, mediated by the evolution of nonlinear dose-response curves, resembles the theoretical framework of Wright (*49*) and Rendel (*50*), describing how gene activity links to phenotypes through a single threshold. Fates of cells close to the threshold exhibit high sensitivity to mutational inputs and evolve fast. Conversely, cells at the low and high end of the curve exhibit low variance in response to a given perturbation, that is, canalisation (*51*) or robustness (*52*). Accordingly, we propose a mechanistic model of phenotypic evolution in which the developmental sensitivity of precursor cells channels phenotypic variation during the evolutionary process. We expect that the distinct positions of the cells in the Wnt dose-responses curve of *Caenorhabditis* and *Oscheius* are due to quantitative changes in the Wnt signalling pathway or in factors acting in parallel to it. In this scenario, reduced Wnt signalling in *Oscheius* vulva precursor cells would position P4.p at the fading end of the activity gradient, making it the most developmentally sensitive cell, while P3.p would remain outside the sensitive zone, rendering it insensitive to incoming variation. In this model, there is no discontinuity between the situations in *Caenorhabditis* and *Oscheius,* with *C. briggsae* located at a crucial transition point where P4.p starts to be affected (Fig. 3). The strength of this model is that a single dose-response curve can account for evolution across different scales, within each clade as well as between the two clades.

Altogether, our work shows how the evolution of the variational properties of cell fate decisions explains the evolution of evolutionary rates for different traits and can be explained by a simple shift of cells in a dose-response curve of a signalling pathway. Developmental bias therefore emerges as a central mechanism connecting micro- and macro-evolutionary processes. These results emphasise the importance of revealing the evolving structure of the genotype-phenotype map to understand phenotypic variability and its evolution across evolutionary time scales.

## Supporting information

Supplementary tables (others in main pdf)

## Acknowledgments

We thank John Wang, Takao Inoue, Erik Andersen, Yen-Ping Hsueh, Walter Sudhaus, David Fitch for strains, Karin Kiontke for help in *Oscheius* species collection and delineation, Hervé Gendrot for plate pouring. We are grateful to Pierre de Villemereuil, Katie Pelletier, Marie Marcaillou, Henrique Teotónio, François Mallard and Luke Noble for discussions and/or providing useful comments on the manuscript. Some strains were provided by the CGC, which is funded by NIH Office of Research Infrastructure Programs (P40 OD010440). This research was funded in part, by the Wellcome Trust Grants 220540/Z/20/A. For the purpose of Open Access, the author has applied a CC BY public copyright licence to any Author Accepted Manuscript version arising from this submission.

## Funding

Agence Nationale de la Recherche (France) 18-CE13-0006-01 (MAF, CB)

Agence Nationale de la Recherche (France) 22-CE13-0005-01 (MAF, CB)

Marie Skłodowska Curie Training Grant (European Union) 751530-EvoCellFate (JPO)

Marie Skłodowska-Curie Actions (European Union) 101110355-EvoBias (JPO)

Wellcome Trust (United Kingdom) 220540/Z/20/A (MB, LS, PGR)

## Author contributions

Conceptualization: JPO, MAF, CBr Investigation: JPO, CBo, NF, MAF, PGR, LS Visualization: JPO, MAF Funding acquisition: MAF, CBr, MB Supervision: MAF, CBr, MB Writing – original draft: JPO, MAF Writing – review & editing: CBr, LS

## Data and materials availability

All raw data are available in the supplementary materials. The data analysis code and plots of the posterior distributions of the models are available at https://github.com/Picao-Osorio/Evolution-Developmental-Bias/tree/main. Links to genome assemblies used in the phylogenetic analyses are provided in table S2.

## Supplementary Materials

Materials and Methods

Figs. S1 to S17

Tables S1 to S9

References (54–105)

Data S1 to S7

## Materials and Methods

### Overall design of the study

We phenotyped vulval precursor cell (VPC) fates (Fig. S1) at three evolutionary scales in the same environment: among species (*V_R_*, rate among species genetic variance), within species (*V_G_*, standing genetic variance), and in *de novo* Random Mutant Lines (RML) together with their respective ancestral control lines (*V_M_*, mutational variance) (Fig. S2). The *V_G_* and *V_M_* datasets were acquired in parallel in a block design.

To determine the rate of phenotypic evolution in two clades, we first determined vulval fates in available species of the *Caenorhabditis* and *Oscheius* genera (53 *Caenorhabditis* species and all available 24 *Oscheius* species), for which we built a phylogeny from genomic and transcriptomic data. For many *Caenorhabditis* and *Oscheius* species, only a single wild isolate is available and we used the reference strain of each species.

To estimate the amount of heritable phenotypic variation within species (standing genetic variance, *V_G_*, expressed in square units of phenotype), we assayed wild isolates of four species. These four species were chosen as representative of the two genera because of their androdioecious reproductive system (hermaphrodites and males) and large available collections of wild isolates. We chose 50 isogenic wild isolates of *C. elegans, C. briggsae* and *O. tipulae* based on their broad geographic distributions and genetic distances (*54–56*); and all 16 available *O. onirici* wild isolates. The isogenic wild isolates were previously established by self-fertilisation of single individuals isolated from the wild.

To estimate the amount of heritable phenotypic variation resulting from random *de novo* mutation (mutational variance, *V_M_*,), we first generated random mutation lines (see below and Fig. S2A) in two wild isolate background of each of the four species. We assayed them in parallel with the wild isolates to assess the mutationally accessible phenotypic spectrum for the six cells P3.p to P8.p, in these two clades (Maynard Smith et al. 1985; Lynch and Walsh 1998).

The data were analyzed using a quantitative genetic framework accounting for non-genetic factors (see below). We then used a similar quantitative genetic framework to estimate the evolutionary rate among species within each genus. In this way, we account for environmental variance to estimate a “rate among species genetic variance” *V_R_* (Housworth et al. 2004; Houle et al. 2017).

For readers unfamiliar with the quantitative genetic framework, consider these variances as measures of phenotypic variation for each trait (the fate of each cell) at three evolutionary timescales: in response to random mutation (*M*), and in intraspecific (*G*) and interspecific (*R*) variation. The variance in fates of the six cells are directly comparable on the same scale.

Taken together, we present here findings based on the phenotypic analysis of over 80,000 individual animals, grown under identical laboratory conditions and examined using Nomarski microscopy. The raw data for each individual is shown in Data S1 and S6. We represent results on the scale of phenotypes in Fig. 2 and of estimated variances in Fig. 3. Supplementary figures display various analyses of the data.

### Nematode strains and culture

All nematode species and strains were maintained in standard laboratory conditions at 20°C on NGM (nematode growth medium) plates with *Escherichia coli* OP50 as a food source (*57*). A complete list of strains used in this study is provided in Table S1.

### Phylogeny

To generate a phylogeny for Rhabditina (clade V) (Fig. S3), we used BUSCO 5.2.2 (*58*) to analyze the genomes of 23 nematode clade V species and a clade IV outgroup (first sheet of Table S2). We used busco2fasta (available at https://github.com/lstevens17/busco2fasta) to identify and extract the protein sequences of 464 BUSCO genes (337,234 sites) that were single-copy in at least 20 (85%) of the 23 genomes. We aligned the protein sequences using MAFFT v7.49 (*59*) and used IQ-TREE 2.2.0.3 (*60*) to infer a gene tree for each BUSCO gene, allowing the best-fitting substitution model to be automatically selected (*61*). We provided the resulting gene trees to ASTRAL-III 5.7.8 (*62*) to infer the species tree. We concatenated the alignments of each protein sequence into a supermatrix using catfasta2phyml v1.1.0 (available at https://github.com/nylander/catfasta2phyml) and estimated branch lengths in substitutions per site for the ASTRAL-III species tree using IQ-TREE under the LG substitution model (*63*) with gamma-distributed rate variation among sites.

To infer a phylogeny for *Caenorhabditis* and *Oscheius* (Fig. 1A, S4), we ran BUSCO 5.2.2 (*58*) on the genomes and transcriptomes of 56 *Caenorhabditis* species and 25 *Oscheius* species (second sheet in Table S2). We used busco2fasta to identify and extract the protein sequences of 560 BUSCO genes (477,102 sites) that were single copy in at least 74 (91%) of the 81 genomes. We inferred a species tree as previously described: we aligned the protein sequences using MAFFT, inferred gene trees using IQ-TREE, species tree using ASTRAL-III, and branch lengths in substitutions per site using catfast2phyml and IQ-TREE, as previously described.

This considerably extends the *Caenorhabditis* phylogeny (*64, 65*) and of the *Oscheius* genus (*26*).

The set of *Oscheius* species appears somewhat less diverse at this genomic level than the set of *Caenorhabditis* species, albeit not greatly (Fig. S4).

### Dating and chronogram

We time calibrated the phylogenies in order to estimate phenotypic evolutionary rate (*V_R_*, see below). We calibrated the chronogram of *Caenorhabditis* and *Oscheius* species (Fig. 1C, S5, S6) by constraining the tree with three *Caenorhabditis* divergence times previously estimated (*66*) with the following corrections: a 14-day generation time estimated from a natural population (Wei, Félix et al. in preparation) and a corrected mutation rate (*μ*) of 2.19×10^-9^ per site, per generation. This mutation rate was obtained by the weighted average of four recent studies that estimated *C. elegans* (N2) mutation rate using spontaneous mutation accumulation lines (MAL): 1.84 ×10^-9^ with 10 MALs (*67*), 2.33×10^-9^ with 32 MALs (*68*), 2.7×10^-9^ with 10 MALs (*69*), and 1.33×10^-9^ with 7 MALs (*70*). These corrections resulted in the following estimated times of divergence: *Caenorhabditis briggsae* and *Caenorhabditis sinica* between 16.97 to 19.15 million years ago (MYA); *Caenorhabditis tropicalis* and *Caenorhabditis brenneri* between 20.22 to 22.36 MYA; and *Caenorhabditis elegans* and *Caenorhabditis inopinata* between 20.14 and 22.48 MYA. We estimated an ultrametric chronogram using the semiparametric method penalized likelihood (PL) with the Truncated-Newton (TN) algorithm for optimising the likelihood of branch lengths and an additive penalty implemented in the program r8s v.1.7 (*71*) (method=pl; algorithm=tn; penalty=add). The PL method models different substitution rates among branches with a penalty for rates changing abruptly from nearby branches that is determined by a ‘smoothing parameter’. We estimated the best smoothing parameter as **1** using cross-validation of a wide range of smooth parameters from 0.01 to 10,000 (crossv = yes; cvstart = −2; cvinc = 0.5; cvnum = 15). This low smoothing value indicates a strong deviation from a strict molecular clock. Moreover, we used the gradient function to check for multiple solutions using different initial starting points (checkGradient=yes, num_time_guesses=3). We obtained an estimated divergence time of 160.53 MYA between the *Caenorhabditis* and *Oscheius* genera, 81.58 MYA for the *Oscheius* genus (*O. tipulae* and *O. myriophilus*), and 106.30 MYA for the *Caenorhabditis* genus (*C.* sp. 41 and *C. krikudae*) (see Data S4, Fig. 1 and S5)

We estimated the Rhabditina chronogram (Data S5; see Fig. S3B) in a similar manner with a best smoothing parameter of **0.1**, using the following fixed divergences times from the above analysis:160.53 MYA between the *Caenorhabditis* and *Oscheius* genera; 81.58 MYA for the *O. tipulae* and *O. myriophilus*; 56.47 MYA for the *O. tipulae* and *O. dolichura*; 38.67 MYA for *C. elegans* and *C. japonica*; and 69.48 MYA for *C. japonica* and *C. angaria*.

The set of *Oscheius* species roots somewhat later than the set of *Caenorhabditis* species, between the branching dates of *C. parvicauda* and *C. plicata* (Fig. S5).

### Generation of mutation lines

We generated eight panels of random mutation lines (RML) from hermaphroditic isogenic wild isolates of two *Caenorhabditis* and two *Oscheius* species: JU1200 and PB306 for *C. elegans*, AF16 and PB800 for *C. briggsae*, CEW1 and JU178 for *O. tipulae*, and PS2068 and JU77 for *O. onirici*. The wild isolates AF16, CEW1 and PS2068 were chosen for being reference strains of the respective species (*72–74*). JU1200 was chosen for being the wild isolate closest genetically to the *C. elegans* reference strain N2 (*75*). We did not use the N2 strain because it harbours many known laboratory-adapted polymorphisms (*76*) that could have an impact on our study. The wild isolates PB800, JU178 and JU77 were selected for their genetic and/or phenotypic distance to the respective reference strains (*25, 27, 55, 56*); and PB306 because it was used in a previous study on vulval mutational variance with spontaneous mutation accumulation (MA) lines. The RMLs were generated by chemical mutagenesis (ENU) following the approach of (*77*) (see Fig. S2) to ensure genome-wide mutagenised lines (data not shown). We used N-ethyl-N-nitrosourea (ENU) as mutagen to obtain a broader distribution of molecular lesions compared to ethylmethanesulfonate (EMS) (*77*), and avoid the laborious construction of spontaneous MA lines. Importantly, the molecular mutational distribution is similar between ENU and MA lines (*77*) and our results compare well with those in (*16*) for the effect of spontaneous mutation on VPC fates in *C. elegans* and *C. briggsae* mutation accumulation lines.

Briefly, P0 larvae undergoing germline proliferation and differentiation at the 3^rd^/4^th^ instar were incubated with 5 mM ENU for four hours at room temperature in a rotary shaker. For the *Oscheius* strains before adding the mutagen, nematodes were incubated in 0.25% SDS for 2 minutes to increase the tissue penetrance (*78*) to the mutagen. Several mutagenised P0s were placed on each plate and F1 animals were screened and selected for the dominant twitching phenotype (*unc-22*[-/+]) in 1% nicotine (*79*). This F1 selection was conducted to ensure that gametes had been effectively mutagenised since *unc-22* is a very long protein-coding gene in *C. elegans* as well as the other species and its mutant phenotype is highly specific. Next, F2 individuals were counter-selected for the twitching phenotype to generate *unc-22*[+], isogenised by 10 generations of selfing hermaphrodites and cryopreserved using standard methods (*57*). At each generation, a randomly early larva was picked to establish the next generation, thus avoiding selection biases. Importantly, for a given ancestral genotype, only RMLs derived from different P0 plates were phenotyped to guarantee their genetic lineage independence.

Before the mutagenesis, we generated two Ancestral Pseudo-Lines (APL) for each of the eight ancestral wild isolates (e.g. JU1200-A and JU1200-B). This was accomplished by establishing derived lines of the wild isolates from a single individual, allowed the population to expand and cryopreserved several copies of an APL as before (see below). All RMLs and APLs were generated in the laboratory in Paris.

### Block structure and vulval phenotyping

The nematode phenotyping was conducted by four experimental observers: J.P.-O., N.F. and M.-A.F. in Paris, and C. Bouleau in Nice.

The phenotyping assays to estimate the within species standing genetic variance (*V_G_*) and mutational variance (*V_M_*) were performed together for *V_G_* and *V_M_* in a blind replicated block design by the four experimental observers. The block design accounts for environmental effects within- and among-lines and across experimental phenotyping blocks, and observer bias. All the blocks had control ancestor APLs to account for environmental and non-mutational effects (*3, 80*) (p. 332) (*15*). In the beginning of each block, strains were given a random number for the observers to phenotype strains blindly to the genotype, i.e. wild isolates, RMLs and control APLs. In total, we performed one common pilot block in the two laboratories (P_PILOT and N_Pilot blocks), 10 blocks in Paris (P1 to P10) and 12 blocks in Nice (N1 to N12). Prior to starting the experimental blocks, an intense training session on vulval phenotyping was conducted in Paris to establish uniformity in ascribing vulval cell fates. Next, the pilot block was conducted to ensure phenotyping consistency among observers, where the same RMLs and APLs from JU1200 (*C. elegans*) and CEW1 (*O. tipulae*) were phenotyped by the four observers. In the next blocks, RMLs and wild isolates were randomly allocated to blocks in the two laboratories. In the P blocks, the three observers randomly phenotyped strains/replicates; while in the N blocks, all strains and replicates were phenotyped by C. Bouleau. A block consisted of strains that were thawed together from APLs and RMLs of a given ancestor/species, and wild isolates of the same species (from the 3^rd^ block onwards), cultured in the same batches of plates and phenotyped in sequential days. The mode value of APLs, RMLs and Wild Isolates per block for a given species was two, four and three, respectively. Most blocks (P1-P8, N1-6/11) had the four species; and only two blocks had species from one genus. Detailed information on the block structure is in Table S5.

Nematode culturing and phenotyping of each block was conducted as in (*16*) but with two replicates (see Fig. S2B). Briefly, a single individual from each thawed line (G0) was arbitrarily selected to initiate the experimental population in order to eliminate potential genetic variation. After two generations of population growth (G1-2), 30-50 gravid adult hermaphrodites per line were cleaned of microbial contaminants by hypochlorite treatment (*57*). Following two discrete generations of culture expansion, 15-30 adult hermaphrodites (G4) were transferred to a new plate to lay the offspring to be phenotyped for replicate A. When the majority of their progeny had reached the L4 stage (G5; after 2-6 days depending on the line, species or genus), vulval cell fates were scored on early to mid L4 larval stages (see below). In parallel, nematode cultures were maintained in standard growing conditions for 3 discrete generations (G4-6’) by agar chunking, then 15-30 adult hermaphrodites at G7’ were transferred to a new plate to lay the offspring to be phenotyped for replicate B (G8) as before. In sum, the control APLs, RMLs and the wild isolates were treated in the same way and blindly, underwent approximately 5 and 8 generations for the two replicates, respectively, between thawing and phenotyping on NGM plates at 20°C and low density. To avoid morphological selection bias for the phenotyped specimens, nematodes were washed from the plate with M9 buffer, spun-down, mounted onto an agar pad, and the first 50 early to mid L4 larvae were phenotyped with Nomarski microscopy with a 100x objective. The pads were made of 6% noble agar and 3 mM of sodium azide (anaesthetic) (*81*) with a Vinyl Record (*82*) to keep the nematodes straight to facilitate the low-throughput phenotyping. Standard criteria were used to infer vulval cell fates based on the arrangement, topology, size and number of cells (*83–85*). P(5–7).p adopt vulval fates and divide to form the adult vulva. The three other cells are competent to form the vulva but normally fuse with the epidermal syncytium hyp7 either before division (L2 stage) or after division (L3 stage). In *Oscheius tipulae*, the most common cell lineage is two rounds of divisions (four cells, “SSSS”) but variants with 3, 2 or 1 cell are also seen. Here we simplify the dataset for *Oscheius* and code the phenotype as binary: two divisions versus other fates. For further information on the rhabditid family and outgroup, see (*26, 40*).

The culturing and phenotyping of *Caenorhabditis* and *Oscheius* species were performed independently by two observers. In general, each species was phenotyped in two consecutive replicates of 50 specimens each. Detailed information on the number of strains and specimens phenotyped are in Tables S3, S6 and S7 below. Detailed phenotyping of all individuals is available in Data S1 and S6.

### Quantitative genetic analysis

The analysis of the phenotyping data was performed using R (v. 4.2.3) (*86*) in RStudio (v. 1.4.1106) (*87*). To estimate quantitative genetic (QG) parameters of the Pn.p vulva precursor cells (VPC) at the three evolutionary scales (*V_R_*,*V_G_* and *V_M_*), we used the phenotypic data among lines to run Bayesian generalised linear mixed models, using the R package MCMCglmm (v. 2.36) (*88*). We focus here on univariate models for each cell. The univariate models were constructed following the course notes of MCMCglmm and the tutorials of the R package QGglmm (*89*). We considered the six vulval traits (P3.p to P8.p) to follow a binomial distribution, where the most common fate for a given VPC was assigned a value of 1 and all others a value of 0. That is, in *Caenorhabditis* species the most common division fate for P3.p, P4.p and P8.p is one division (fate ‘SS’), while in *Oscheius* species it is two divisions (fate ‘SSSS’). For the central VPCs P5.p to P7.p, we considered their canonical 2°, 1°, 2° fates, respectively, as ‘wt’ with a value of 1. Given the binomial distribution of all traits, we used the threshold model (*49*) with a probit link-function that has a better mixing of the Markov Chain Monte Carlo (MCMC) chains (*90*). To ensure large MCMC effective sample sizes of the posterior distribution, we chose 5,010,000 and 1,260,000 iterations, and 2,000 and 500 thinning intervals for the analyses of *V_R_* and *V_G_* _& *M*_, respectively. This resulted in minimum samples sizes of genetic variance in the liability scale of 2,302 for *V_R_*, 2,244 for *V_G_*, and 2,194 for *V_M_*. In all models, burn-in was set to 10,000 iterations and convergence was checked by visual inspection of the posterior distributions and by the Heidelberg and Welch (*91*) convergence test implemented in the coda R package (v. 0.19-4.1) (*92*). Furthermore, to contribute for model convergence, latent variables were truncated (trunc=TRUE) to prevent under/overflow (-/+7) of the models. The posterior distribution of random effects was saved (pr=TRUE) to obtain the best linear unbiased predictions (BLUP) of the nematode strains for each trait, using the R package postMCMCglmm (v. 0.1-2) (*93*). We used non-informative priors: a wide normal distribution on all fixed effects, and a chi-square distribution with one degree of freedom (V = 1, nu = 1000, alpha.mu = 0, alpha.V = 1) on all random effects due to its better inference for binary traits (*94*). The residual variance had to be fixed to 1 because it is not identifiable in binary responses (see MCMCglmm course notes of and QGglmm tutorial). From these models, we extracted the following QG parameters in the liability scale using their posterior distributions: trait mean (z), genetic variance (*V_g_*), phenotypic variance (*V_P_*) and broad-sense heritability (*H^2^* = *V_g_* / *V_P_)*. The trait mean is the overall mean of the fixed effects of the model. The *V_g_* is half the among-line variance because it is obtained from homozygous isogenic lines that include non-additive effects such as dominance (*95*). The phenotypic variance is the sum of the variances from all random effects: *V_G_*, block and replicate effects (*V_Brep_*), residual variance of 1, and variance explained by fixed effects (*96*). These QG parameters were then computed to the observed data scale with the QGglmm package (v. 0.7.34) (*89*), using the binomial error distribution and integrating over the posterior distribution of each parameter. We extracted for all QG parameters the mean, median, posterior mode, and the 95% and 83% credible intervals from their posterior distributions. Significant differences of posterior distributions can be inferred when their 83% credible intervals do not overlap (*97*). Furthermore, to measure the fold-change differences in genetic variance between P3.p and P4.p, we divided their posterior distributions and the resulting ratio displayed in a log10 scale. We did not mean-scale the variance because binomial traits are already in the same scale and there is an intrinsic relationship between mean and variance. To estimate the evolutionary rate among species (*V_R_*), we additionally used the chronogram of each nematode genus to run generalised linear phylogenetic mixed models (*32*) following the guidelines of (*98*). Although this estimate only measures a certain pattern in interspecific variation (*99*), the goal here is to remove the environmental variance and the influence of possible intraspecific polymorphisms by focusing on the part of the variation that is explained by the phylogeny. Briefly, the same logic of the QG animal models was used, but instead of a pedigree relatedness covariance matrix, we used a phylogenetic covariance matrix. This matrix assumes a Brownian motion model of trait evolution where there is phylogenetic inertia and a constant increase in trait variance through evolution. In this way, phenotypic evolutionary rates are estimated as increases of genetic variance per million years (*V_R_*) (*100*). Additionally, phylogenetic heritability (or phylogenetic signal) is the proportion of phenotypic variance explained by the phylogenetic relationships (*H^2^_R_* = *V_R_* / *V_P_*) (*31*) (*99*) instead of pedigree-based heritability. We first generated the inverse phylogenetic covariance matrix using all the nodes and scaled the total branch length, from root to tips, to one (nodes=“ALL”, scale=TRUE with the inverseA::MCMCglmm function). In these models we did not have fixed effects, and had the following random effect: variance among-species dependent on the phylogeny (*V_R1_*), variance among-species not dependent on the phylogeny (*V_NotPhylo_*), variance among species replicates (*V_replicate_*). In this case, *V_P_* = *V_R_* + *V_NotPhylo_* + *V_replicate_* + residual variance of 1. Since the trees were scale in the analyses, all the random factors are in a standardise scale. To obtain the *V_R_* per million years (MY), we divided the *V_R1_* by the total length of the chronogram: 106.30 MY for *Caenorhabditis* and 81.58 MY for *Oscheius*.

To map the trait evolution along each phylogeny (Fig. S6), we obtained the mean trait frequency per species and estimated ancestral states using a Brownian motion model, using the contMap function in the R package Phytools (v. 2.3-0) (*101*).

The standing genetic variance (*V_G_*) is the genetic variance among wild isolates. The mutational variance (*V_M_*) is the genetic variance among random mutant lines (RML). When estimating these variances, we included the variance among control ancestral pseudo-lines (APL) of the respective species to account for uncontrolled non-genetic variation (*3, 80*) (p. 332). For this, we included a fixed effect named *Treatment* for control vs. mutation/wild. We ran models only with the control APLs, the wild isolates and the RMLs, and compared them to the full *V_G_* and *V_M_* models. We observed significant *V_G_* and *V_M_* for P3.p in all *Caenorhabditis* species and isolates, except in the JU1200 *V_M_*; and for P4.p in all *Oscheius* species and isolates (see Figs. S10 and S15). In all *V_M_* models, we ran models with isolates independently, and with the two isolates together for each species. This species *V_M_* was estimated by including an *Ancestral* (e.g. PB306 vs. JU1200) fixed effect that was nested within *Treatment*. In all aforementioned *V_G_* and *V_M_* models, we included the four experimental observers (J.P.-O., N.F. and M.-A.F., and C.B.-M) as fixed effect to take into account observational phenotyping biases. Also, in all these models we had one random effect of block and replicate (*V_Brep_*) that depicted the experimental structure of the phenotyping.

To estimate mutational effects on the directional changes in the trait means of RMLs in respect to their ancestors, we estimated marginal means of the fixed effect *Treatment* (control Vs mutation) using the R package emmeans (v. 1.10.3) (*102*).

All plots were made using the R package ggplot2 (v. 3.5.1) (*103*).

#### Oscheius tipulae egl-20 experiment

The *Oti-egl-20* mutant (*48*) was scored in the same experimental conditions as above, in parallel with the control reference wild-type *O. tipulae* background CEW1.

## Supplementary Text

### Further variation in cell division patterns in the *Oscheius* genus

In *Caenorhabditis*, the fates of P3.p and P4.p are appropriately coded as binary traits, as only rare individuals displayed a fate other than no or one division (Data S1, S6). In *Oscheius tipulae* and *O. onirici*, the canonical 3° fate lineage consists in two divisions. For the sake of simplicity and easy comparison with *Caenorhabditis*, we binarized the data for the analysis by contrasting this canonical fate of two divisions with all others. This removes some further variation within the *Oscheius* genus (fig. S6), which is observed at all evolutionary levels (species, wild isolates, mutation lines) (Data S1, S6). This finding is in line with the alignment of variances at all evolutionary scales.

We note that the occurrence of one or two divisions can be distinguished genetically in *O. tipula*e (*29, 84*) and that the Wnt pathway mostly affects the probability of two versus no divisions in O*. tipulae* reference CEW1 (*48, 84*). The situation is thus more complex than in *Caenorhabditis*. The variation in phenotype space may correspond in part to variation along the same phenotypic axis as the Wnt pathway and in part to variations in cell cycle regulation that may interact with the Wnt pathway (*29, 48, 84, 104*).

### Analysis of the variation for the six Pn.p cells

In this study we determined the fates of six precursor cells (Data S1, S6). In the main text, we focused on the most anterior cells for the sake of simplicity, P3.p and P4.p for bidimensional plots and P(3–5).p in other cases.

When extending the analysis to six cells, the most interesting pattern that can be seen (Fig. S17) is a relatively high variance for P8.p in *Oscheius* at all scales (V_R_, V_G_, V_M_). This finding is again in line with the alignment of variances at all evolutionary scales.

In accordance with our model of Wnt sensitivity (Fig. 4), P8.p is also sensitive to *Oti-egl-20* mutation in *O. tipulae* (Table S10), much more so than the central cells P(5–7).p and more so than in *C. elegans* (where P8.p did not divide in only 2/218 individuals of the *egl-20(hu105)* animals scored in (*43*)). However, in our model of relative sensitivity in the Wnt gradient (Fig. 4), it is not intuitive that P8.p fate should be more variable than the central cells P(5–7).p since it should receive a higher concentration of Wnt. What are the possible reasons for this?

Competence of the Pn.p cells is confered in part by expression of the Hox gene *lin-39* in P(3–8).p from the erly L1 stage However, the different cells quantitatively differ in their competence due to expression of the more Hox gene *mab-5* in P8.p in *C. elegans*, which renders P8.p less competent (*105*). In addition, the gonadal anchor cell is located close to P6.p and emits LIN-3/EGF signaling that maintains competence and vulval fates in P(5–7).p (*28, 42, 48, 104*). An additional Wnt paralog, *mom-2*, is also expressed in the region of P6.p in both *C. elegans* and *O. tipulae* (*44, 48*) and may contribute to P(5–7).p insensitivity. Altogether, this may explain the relative high variance in P8.p in *Oscheius* and does not contradict our model of sensitivity to Wnt signaling.

---

**Figure S1.**
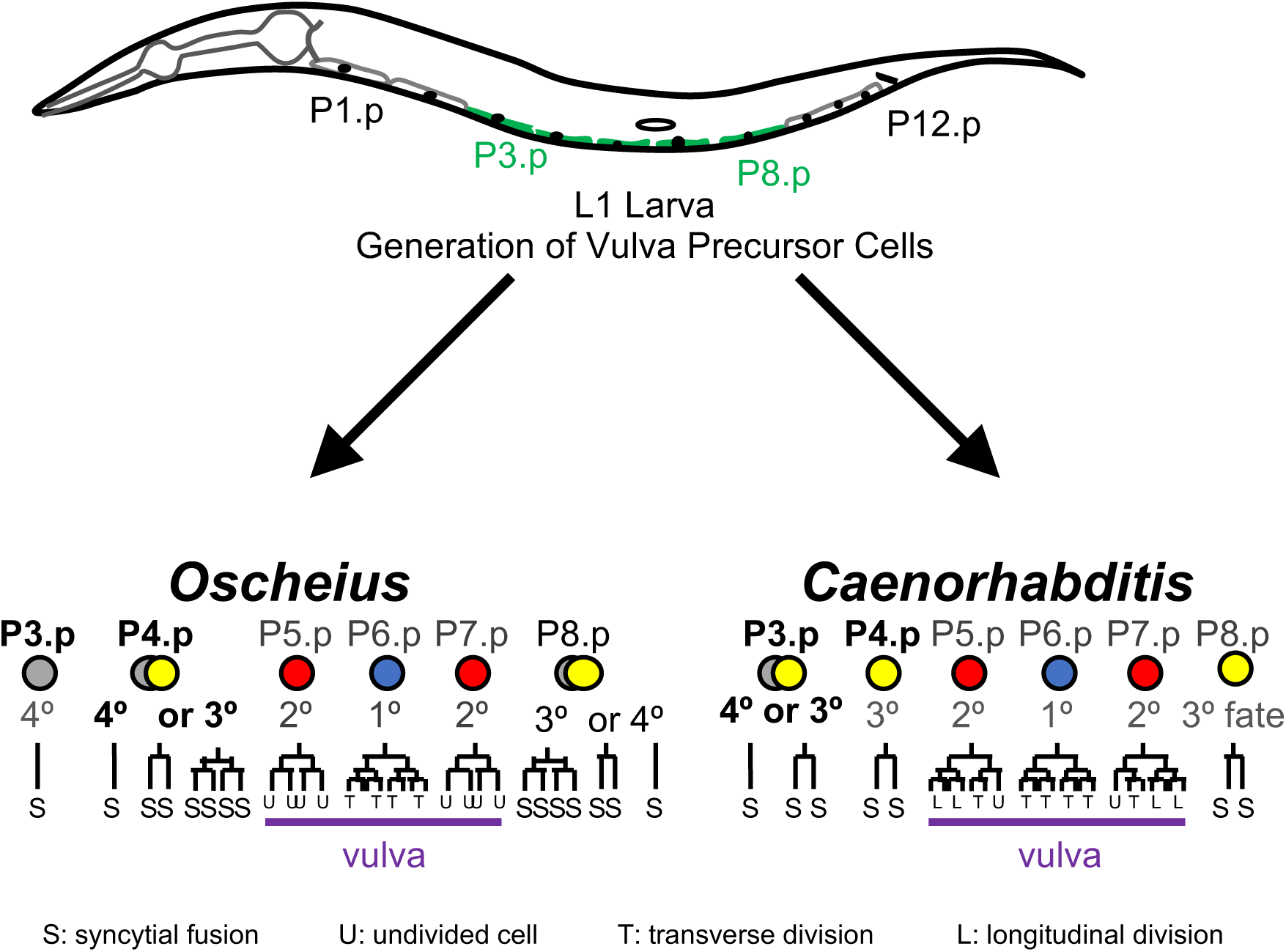
Schematic of vulval development in *Caenorhabditis* and *Oscheius*. The antero-posterior position of the Pn.p cells in the L1 larval stage are shown on the top drawing. These cells are numbered according to their antero-posterior position from n=1 to 12. We consider here the vulva precursor cells P3.p to P8.p (green). The canonical pattern of division of these six cells is shown at the bottom for each genus (*23, 25, 30*). The cell fates are color coded, blue for the central 1° fate induced by LIN-3 signaling from the anchor cell from the somatic gonad, red for the lateral 2°fate induced by Delta-Notch signaling, yellow for the 3° fate (one division), grey (no division). As indicated below the figure, S indicates syncytial epidermal fate (fusion to hyp7). U: undivided grand-daughter of Pn.p cells. T: transverse division in the third round of division. L: longitudinal division in the third round of division.

**Figure S2.**
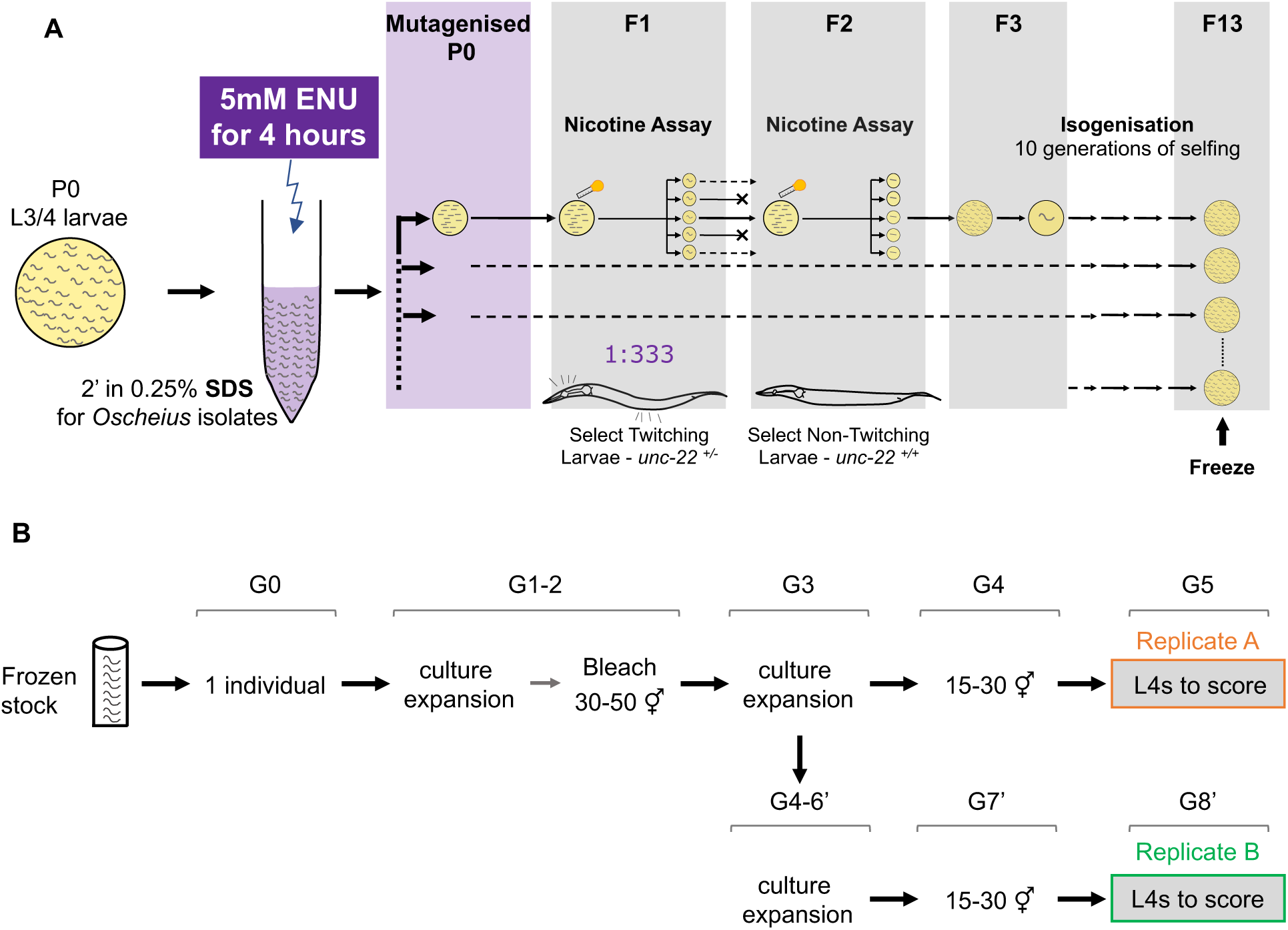
Experimental design for the mutagenesis and the treatment of strains prior to phenotypic scoring. (A) Schematic of the random mutagenesis. Note that in contrast to developmental genetic screens, all lines were kept (if they could be maintained) and scored. See Methods for further details. (B) Schematic of the treatment of the strains prior to phenotypic scoring. Gn denotes the generation.

**Figure S3.**
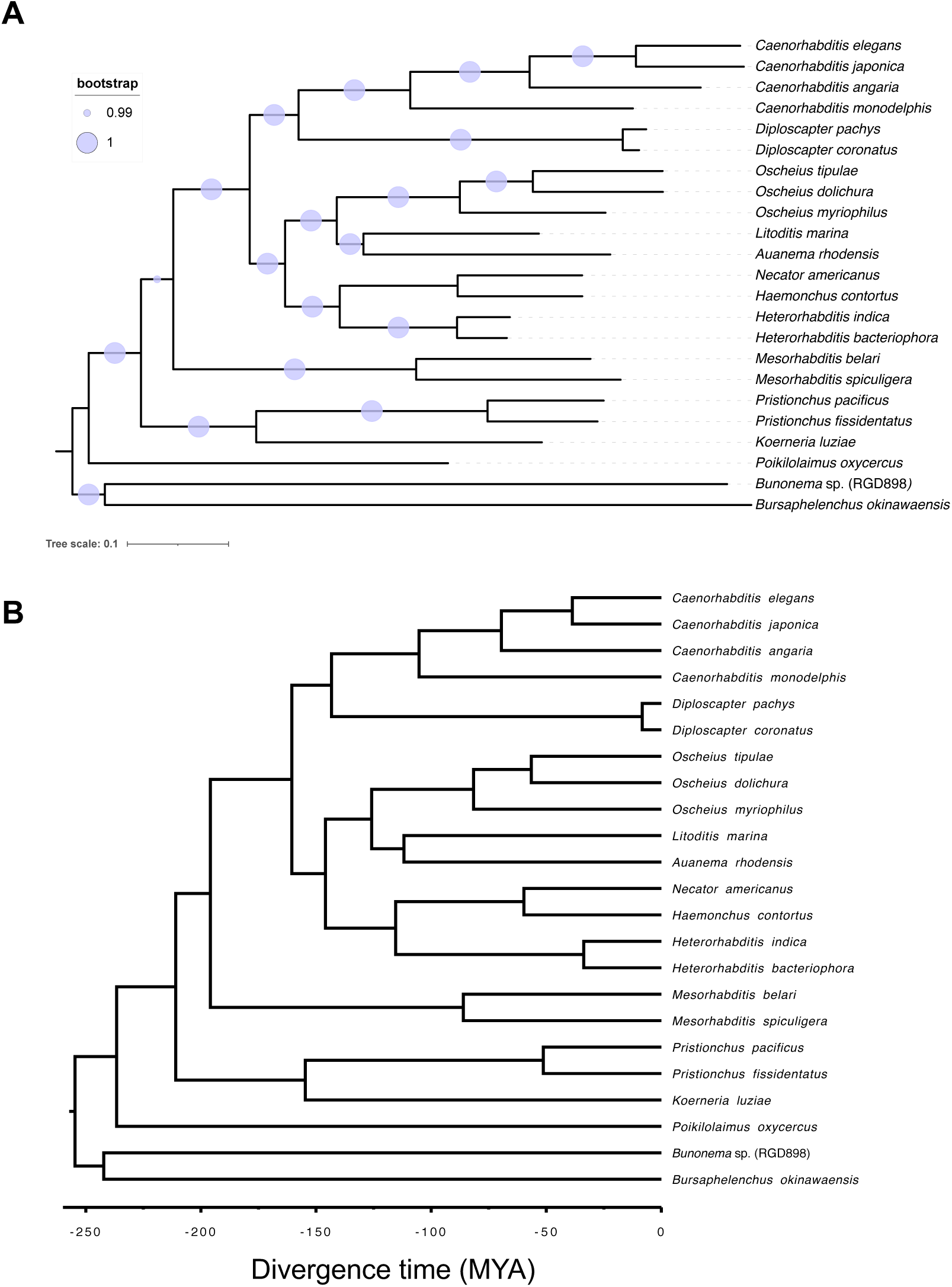
Phylogram and Chronogram of Rhabditina nematodes. (A) Inferred phylogenetic relationships of 23 species of clade V Rhabditina nematodes from 464 BUSCO (benchmarking universal single-copy orthologs) genes using maximum likelihood under the LG substitution model with gamma-distributed rate variation among sites. Branch lengths are in substitutions per site (scale is shown). Bootstraps values are depicted by the size of light blue circles. (B) Inferred chronogram of the same 23 species of clade V Rhabditina nematodes depicting their divergence times in million years ago (MYA).

**Figure S4.**
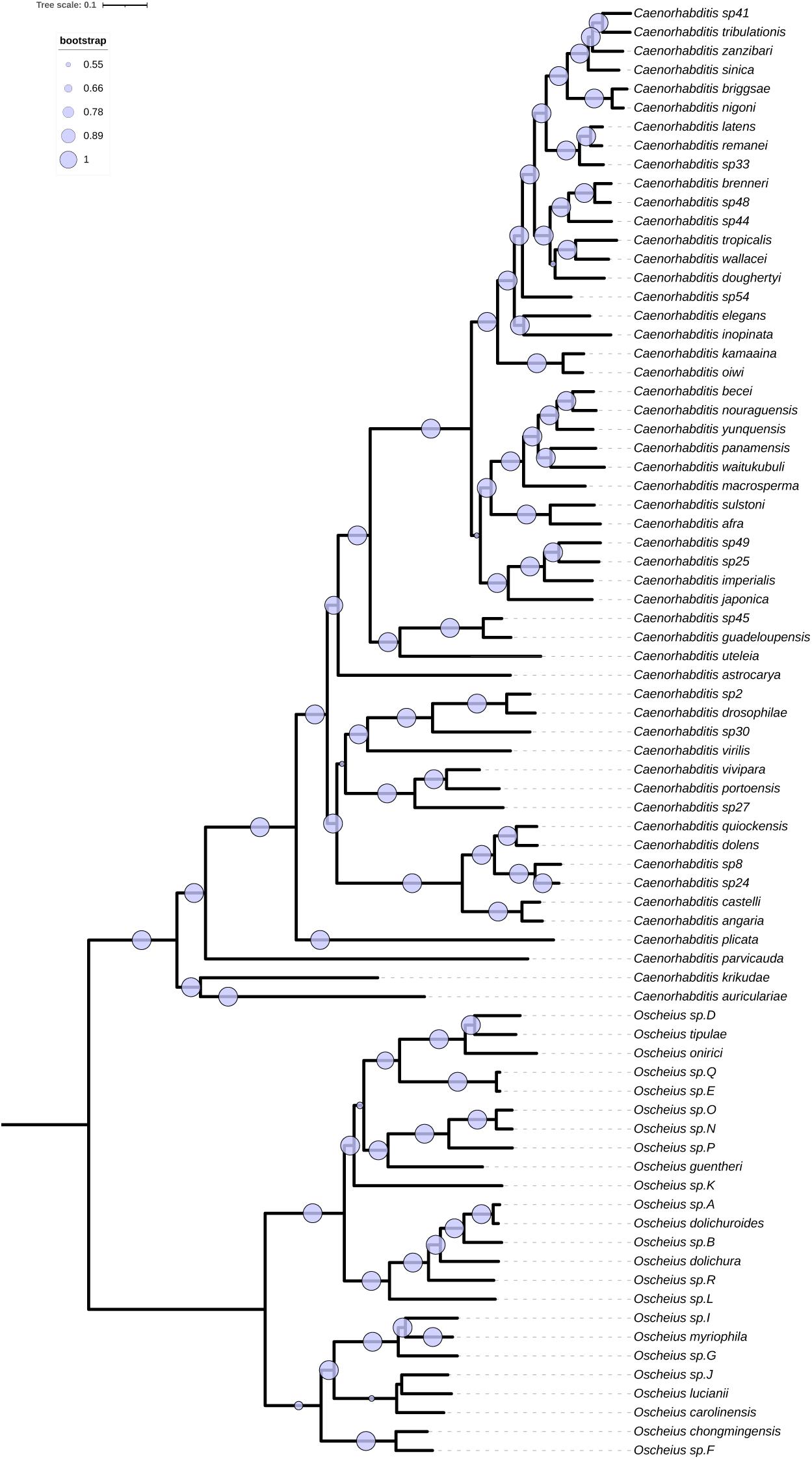
Phylogram of *Caenorhabditis* and *Oscheius* species. Inferred phylogenetic relationships of 56 *Caenorhabditis* and 25 *Oscheius* species from 560 BUSCO genes using maximum likelihood under the LG substitution model with gamma-distributed rate variation among sites. Branch lengths are in substitutions per site (scale is shown). Bootstraps values are depicted by the size of light blue circles

**Figure S5.**
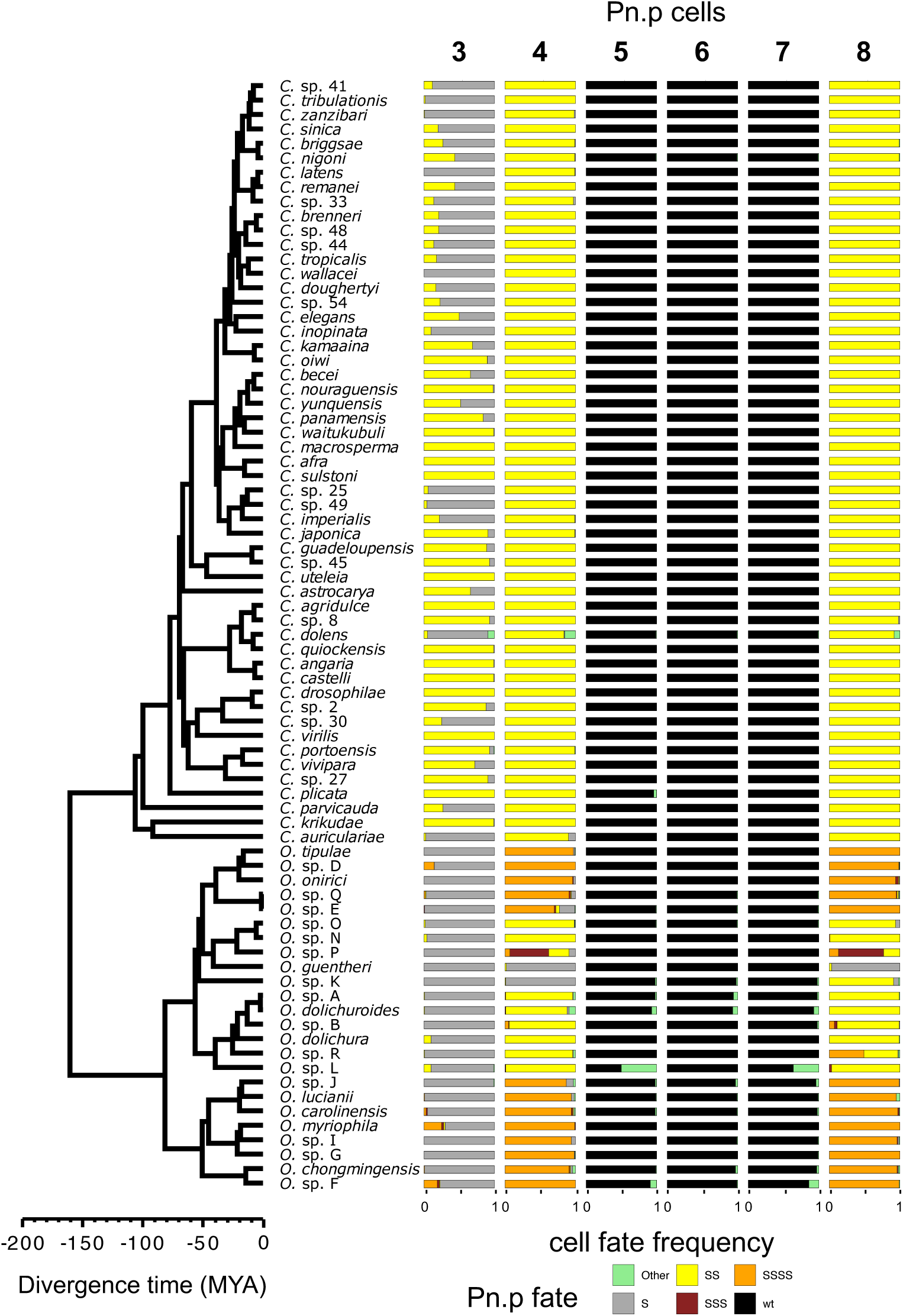
Evolution of vulval cell fates along the chronogram of *Caenorhabditis* and *Oscheius* species. Horizontal bar plots showing the six VPCs cell fates along the chronogram of *Caenorhabditis* and *Oscheius* species: no division (S, grey); one division (SS, yellow); one division of the VPC and division of one daughter (SSS, brown); two divisions (SSSS, orange); 2°, 1°, 2° wild-type vulva fates for P5.p, P6.p and P7.p, respectively (wt, black); other vulval fates (other, green). ‘S’ stands for syncytial fate. Divergence times are in million years ago (MYA).

**Figure S6.**
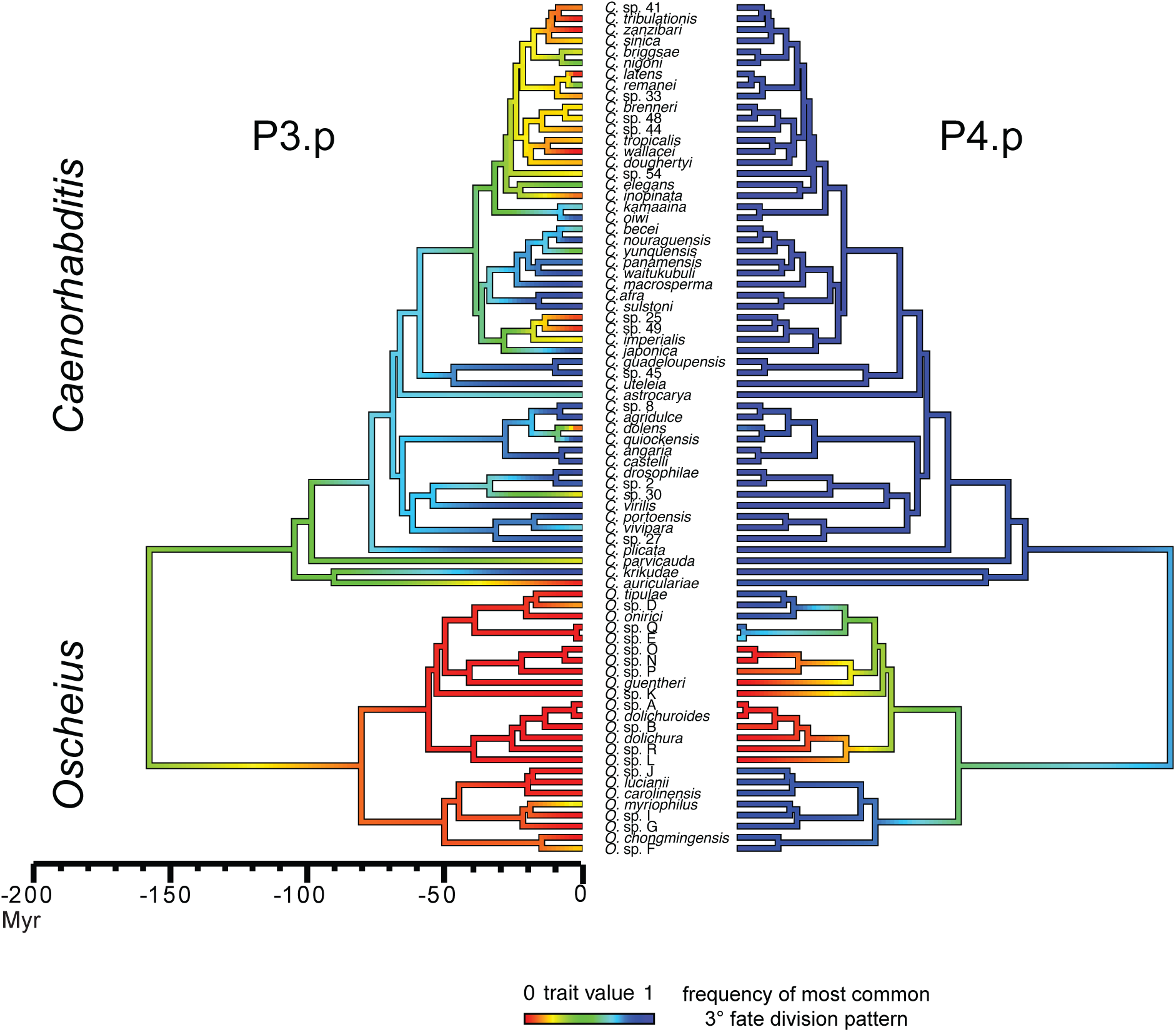
Vulval trait mapping along the phylogeny. Phylogenetic reconstruction of P3.p (left) and P4.p (right) mean division frequency in *Caenorhabditis* and *Oscheius* species represented by changes in colour: 0.0 and 1.0 division frequency are in red and blue, respectively. The trait value refers to the frequency of the most common type of 3° fate division for each genus: one division (cell fate SS) in *Caenorhabditis* and two divisions (cell fate SSSS) in *Oscheius* species. ‘S’ stands for syncytial fate.

**Figure S7.**
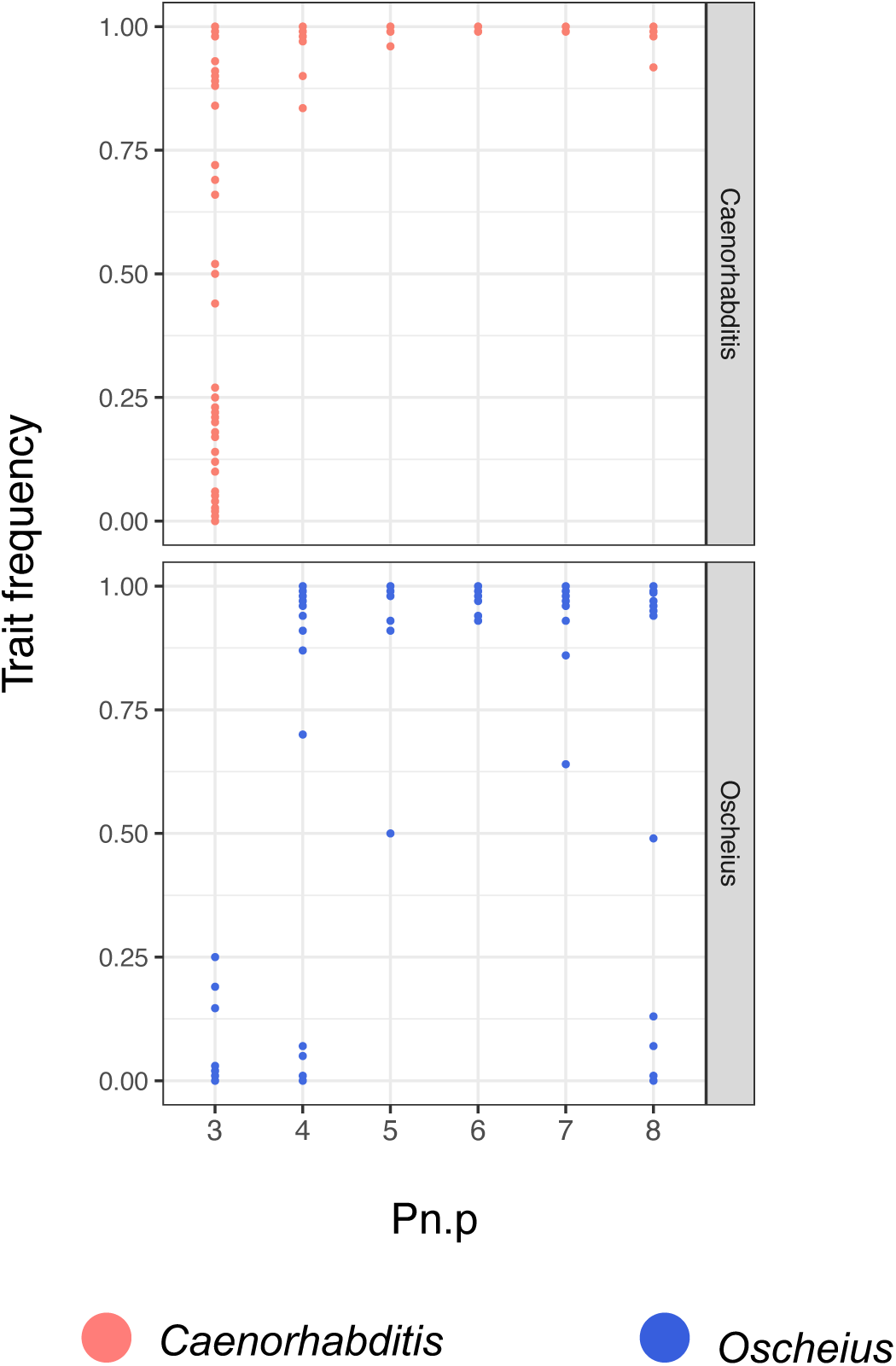
Vulval trait frequency in diverse *Caenorhabditis* and *Oscheius* species. Trait frequency for the six VPCs in species of *Caenorhabditis* (salmon) and *Oscheius* (dark blue). In cases where several isolates were scored, each dot denotes the average trait frequency of the species. In all figures, trait frequency is expressed relative to the most common type of VPC cell fate. For P3.p, P4.p and P8.p, the most common cell fate is one division (cell fate SS) in *Caenorhabditis* and two divisions (cell fate SSSS) in *Oscheius* species. ‘S’ stands for syncytial fate. For P5.p, P6.p and P7.p, the most common cell fate is the 2°, 1°, 2° wild-type vulva fates, respectively. Note that the 2° fated cells in *Oscheius* sp. L included one additional division of one or more of the grand-daughters in about half of the animals (Data S1).

**Figure S8.**
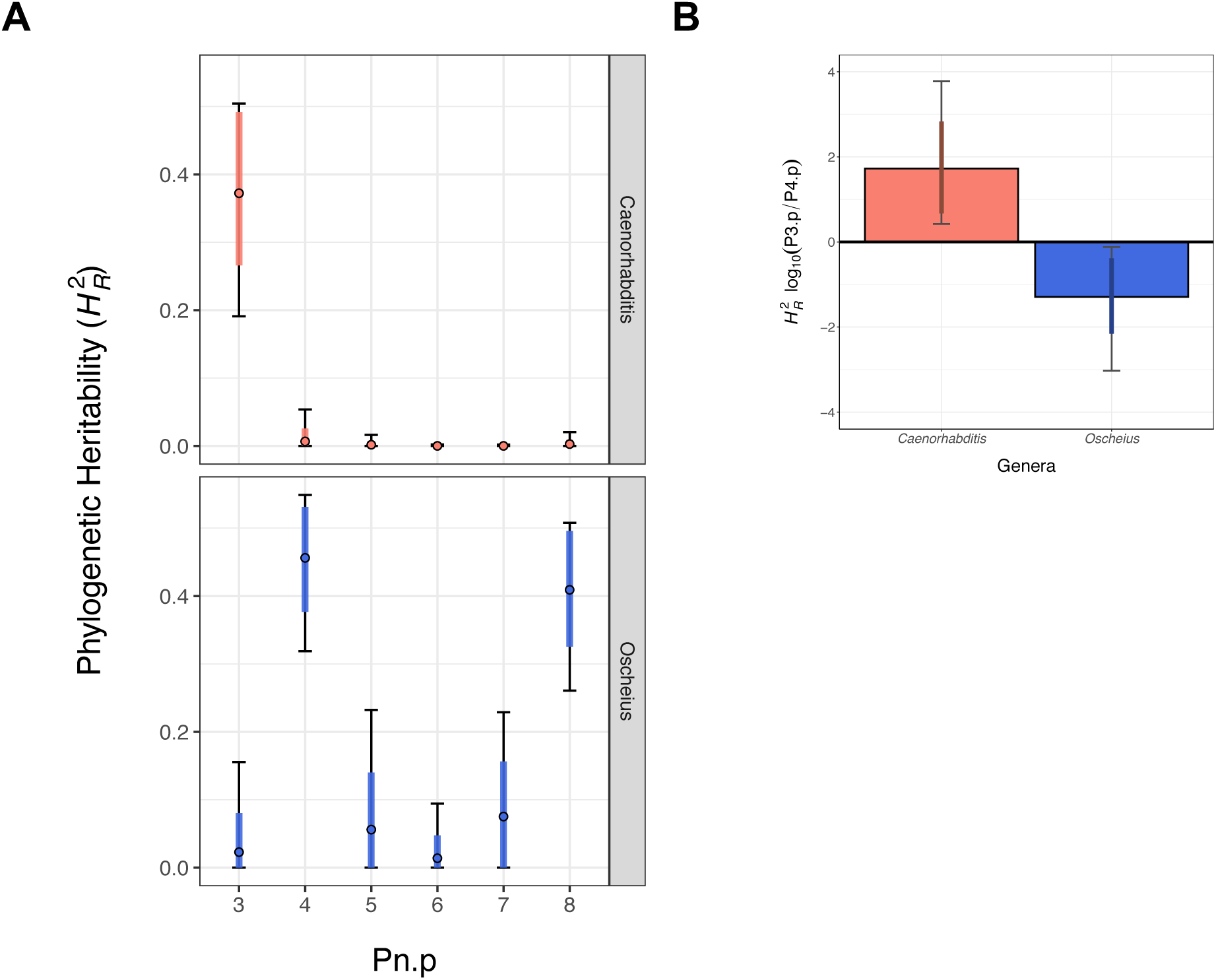
Phylogenetic heritability of vulval cells in the *Caenorhabditis* and *Oscheius* clades. (A) Estimation of the broad-sense phylogenetic heritability (*H*^2^*_R_*) of the six VPCs among species of *Caenorhabditis* (salmon) and *Oscheius* (dark blue). (B) Ten-fold change of the *H*^2^*_R_* ratio between P3.p and P4.p in *Caenorhabditis* (salmon) and *Oscheius* (dark blue). Each estimation displays the median (black circle in A and bar in B), 83% (coloured bar) and 95% (black bar) credible intervals of the posterior distributions in the observed data scale with a minimum effective sample size of 2000.

**Figure S9.**
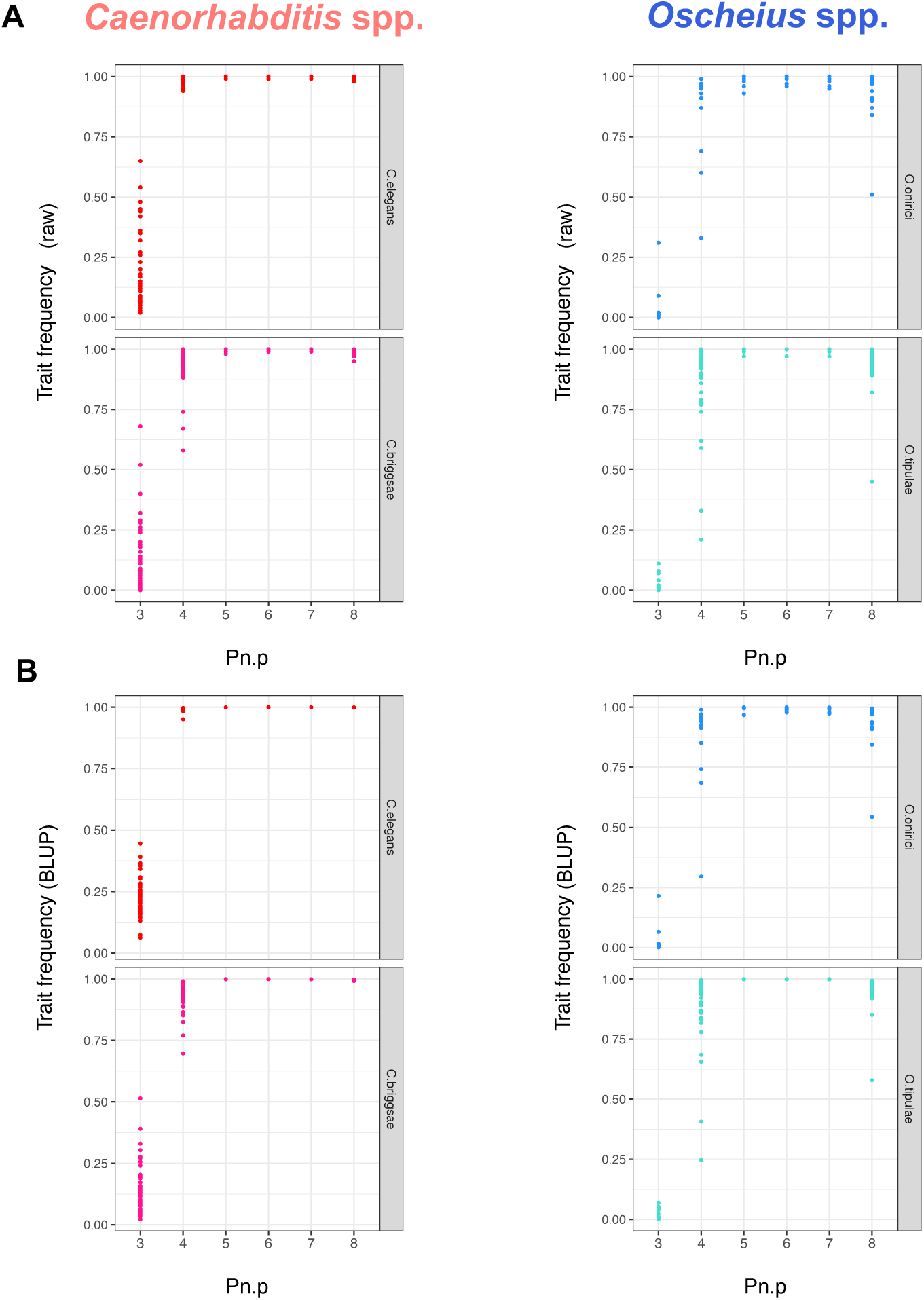
Vulval trait frequency in sets of wild isolates of four *Caenorhabditis* and *Oscheius* species. (A) Trait frequency of the six VPCs in wild isolates of *C. elegans* (red), *C. briggsae* (pink), *O. tipulae* (blue) and *O. onirici* (turquoise). Each dot denotes the average trait frequency of a wild isolate. (B) Best linear unbiased prediction (BLUP) of VPC trait frequency among wild isolates of *C. elegans* (red), *C. briggsae* (pink), *O. tipulae* (blue) and *O. onirici* (turquoise). Each dot represents the BLUP in the observed data scale of a wild isolate that is corrected for the block and observer structure, and the intrastrain variation of two ancestral pseudo-line controls for each species. Trait frequency is expressed relative to the most common type of VPC cell fate as in Figure S7. The raw data for each individual can be found in Data S6.

**Figure S10.**
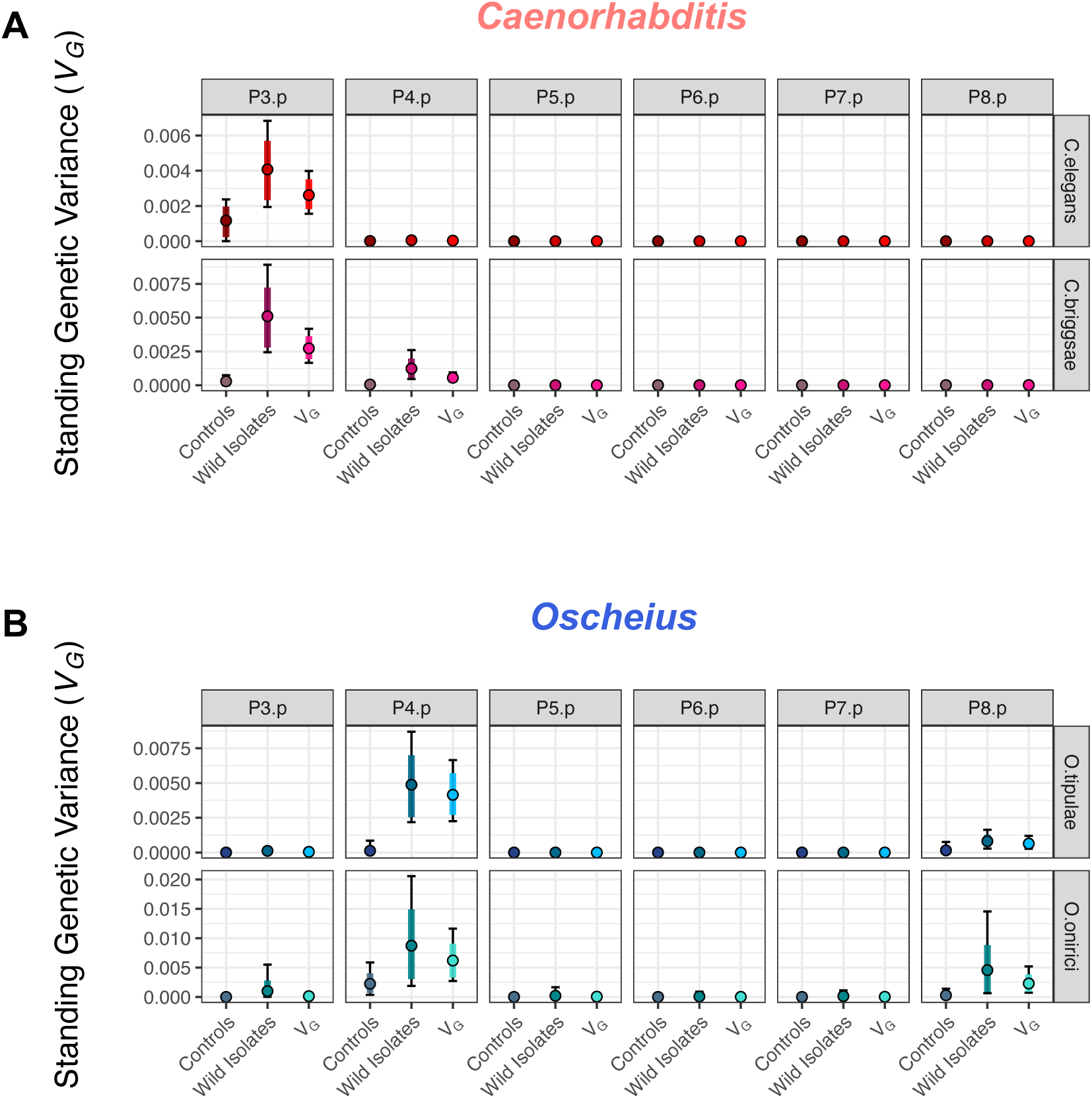
Standing genetic variance of vulval precursor cells within *Caenorhabditis* and *Oscheius* species. Estimation of the standing genetic variances (*V_G_*) of the six VPCs in (**A**) *C. elegans* (red), *C. briggsae* (pink), and (**B**) *O. tipulae* (blue) and *O. onirici* (turquoise). For each VPC are shown: left, intrastrain variation of two ancestral pseudo-line controls; middle, variation among wild isolates; right, full *V_G_* model with a fixed effect named *Treatment* for control vs. wild isolates. Each estimation displays the median (black circle), 83% (coloured bar) and 95% (black bar) credible intervals of the posterior distributions of the genetic variances in the observed data scale. Each posterior distribution has a minimum effective sample size of 2000. The *V_G_* scale is adjusted to each species.

**Figure S11.**
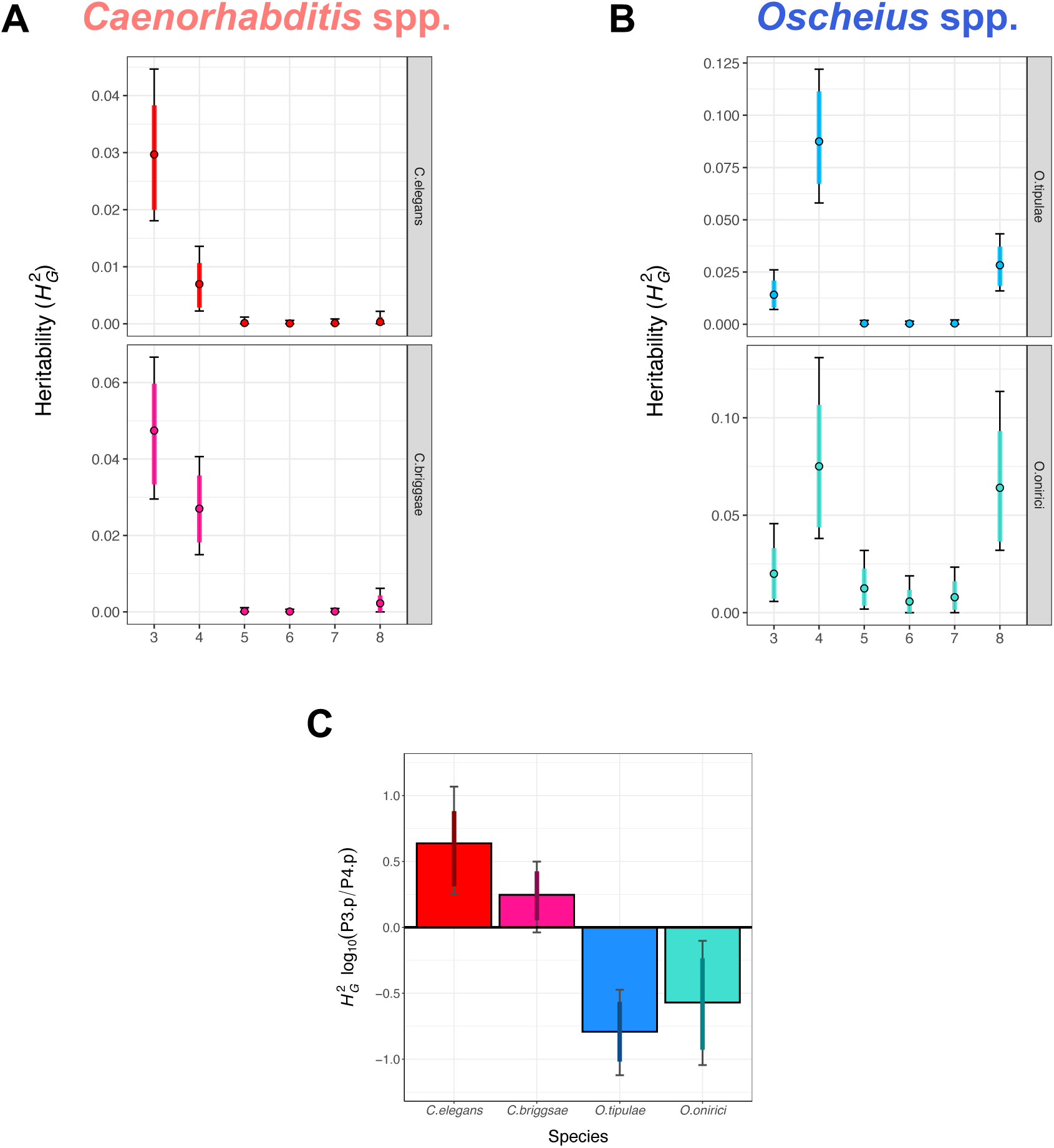
Heritability of vulval precursor cell fates within each *Caenorhabditis* or *Oscheius* species. Estimation of the broad-sense heritability (*H*^2^*_G_*) of the six VPCs in (**A**) *C. elegans* (red), *C. briggsae* (pink), and (**B**) *O. tipulae* (blue) and *O. onirici* (turquoise). The *H*^2^*_G_* scale is adjusted to each species. (C) Ten-fold change of the *H*^2^*_G_* ratio between P3.p and P4.p in *C. elegans* (red), *C. briggsae* (pink), *O. tipulae* (blue) and *O. onirici* (turquoise). Each estimation displays the median (black circle in A and B, bar in C), 83% (coloured bar) and 95% (black bar) credible intervals of the posterior distributions in the observed data scale with a minimum effective sample size of 2000.

**Figure S12.**
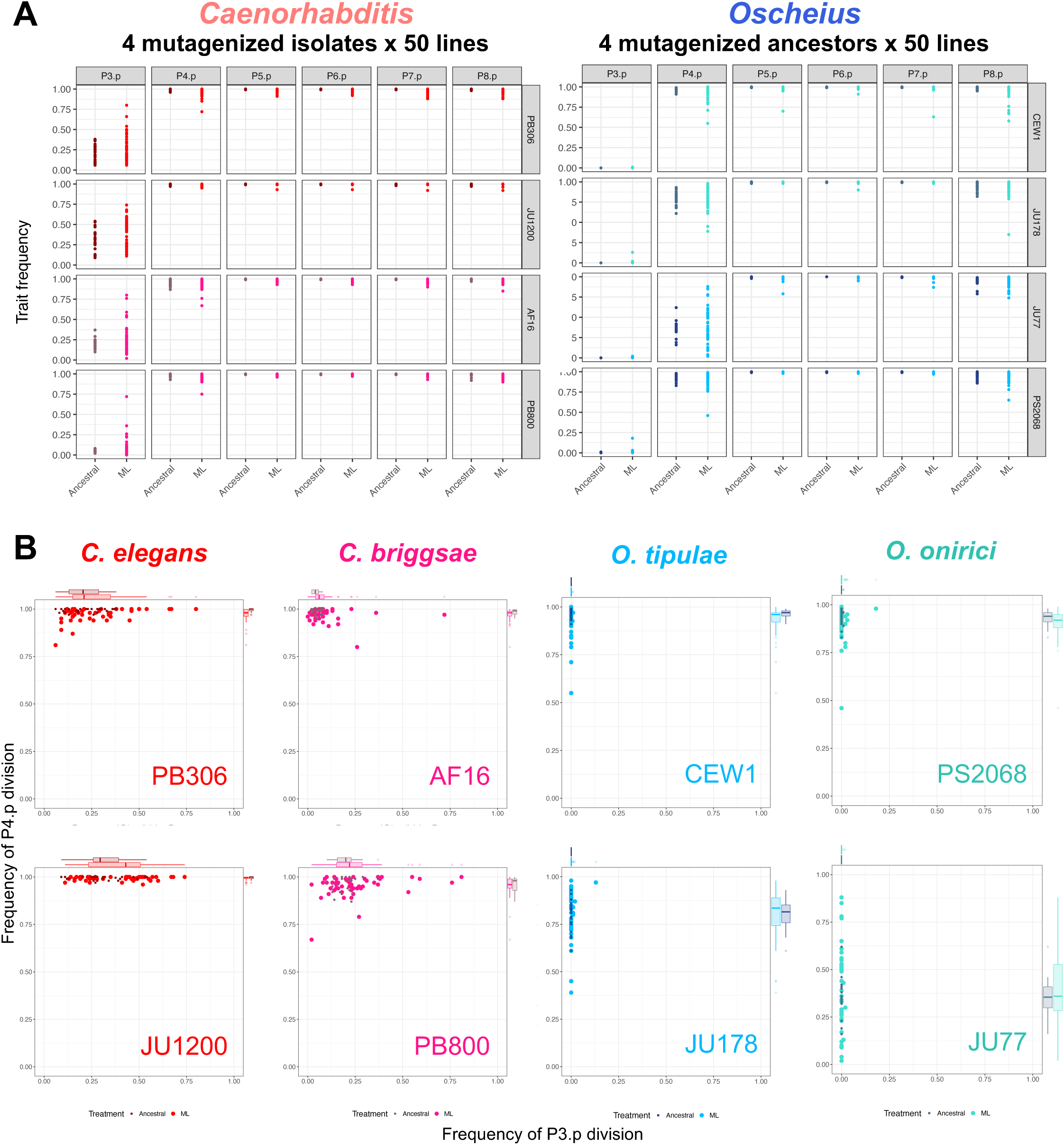
Vulval trait distribution generated from *de novo* mutation in *Caenorhabditis* and *Oscheius* species. (A) Raw data of trait frequency of the six VPCs among ancestral pseudo-line (Ancestral, darker colours) and random mutant lines (ML, lighter colours) in two wild isolates of *C. elegans* (PB306 and JU1200, red), *C. briggsae* (AF16 and PB800, pink), *O. tipulae* (CEW1 and JU178, blue) and *O. onirici* (JU77 and PS2068, turquoise). Each dot denotes the average trait frequency of a strain. (B) Raw data of division frequency of P3.p (x-axis) and P4.p (y-axis) upon random mutations in two isolates of *C. elegans* (red), *C. briggsae* (pink), *O. tipulae* (blue) and *O. onirici* (turquoise). The corresponding model estimates, taking into account block and observer effects, are shown in Figure 2. Trait frequency is expressed relative to the most common type of VPC cell fate as in Figure S7. The raw data for each individual can be found in Data S6.

**Figure S13.**
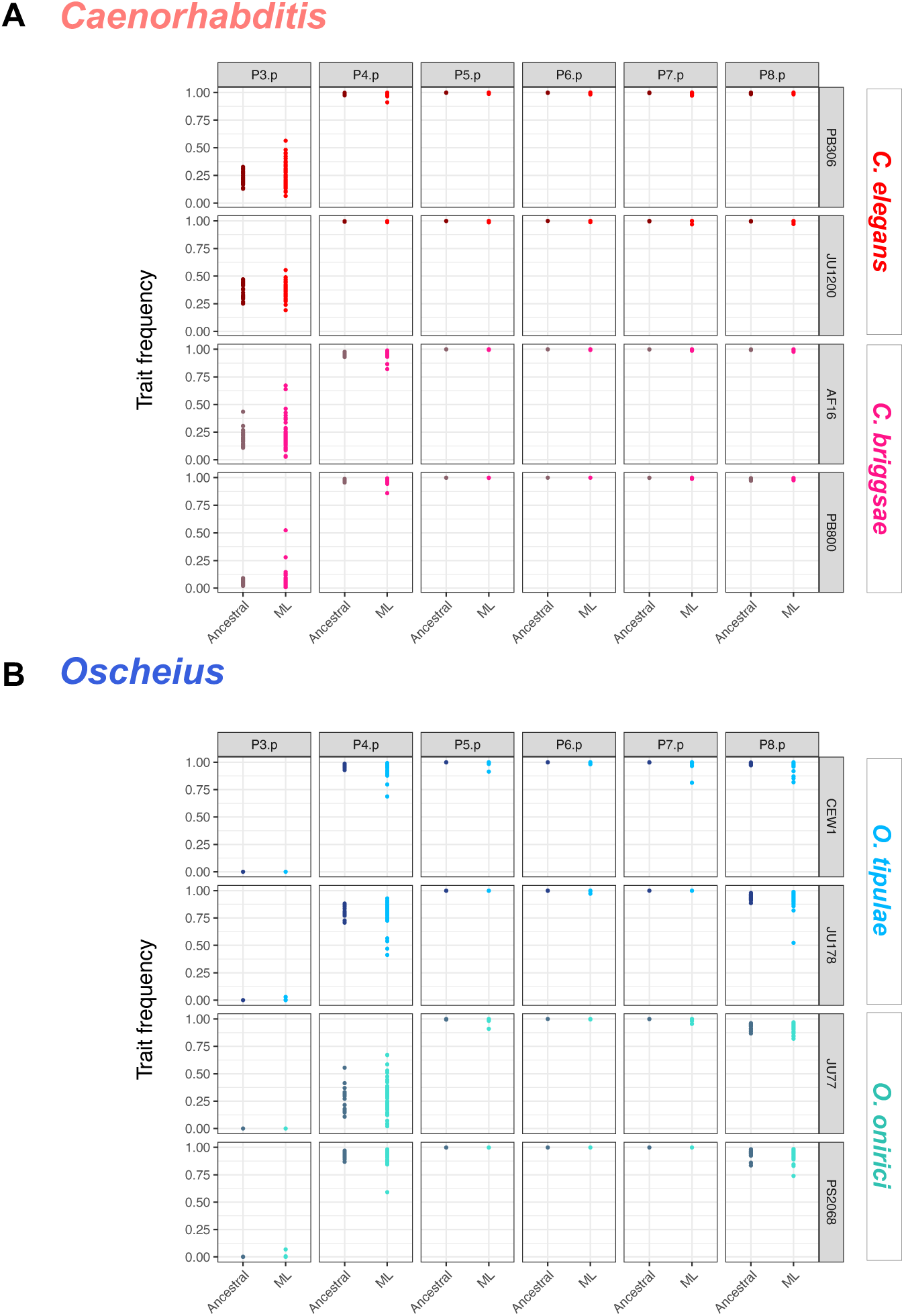
BLUPs of vulval traits upon *de novo* mutation in *Caenorhabditis* and *Oscheius* species. Best linear unbiased prediction (BLUP) of VPC trait frequency in ancestral pseudo-line (Ancestral, darker colours) and random mutant lines (ML, lighter colours) in two wild isolates of (**A**) *C. elegans* (PB306 and JU1200, red), and *C. briggsae* (AF16 and PB800, pink), (**B**) *O. tipulae* (CEW1 and JU178, blue) and *O. onirici* (JU77 and PS2068, turquoise). Each dot represents the BLUP in the observed data scale of strain corrected for the block and observer structure. Trait frequency is expressed relative to the most common type of VPC cell fate as in Figure S6.

**Figure S14.**
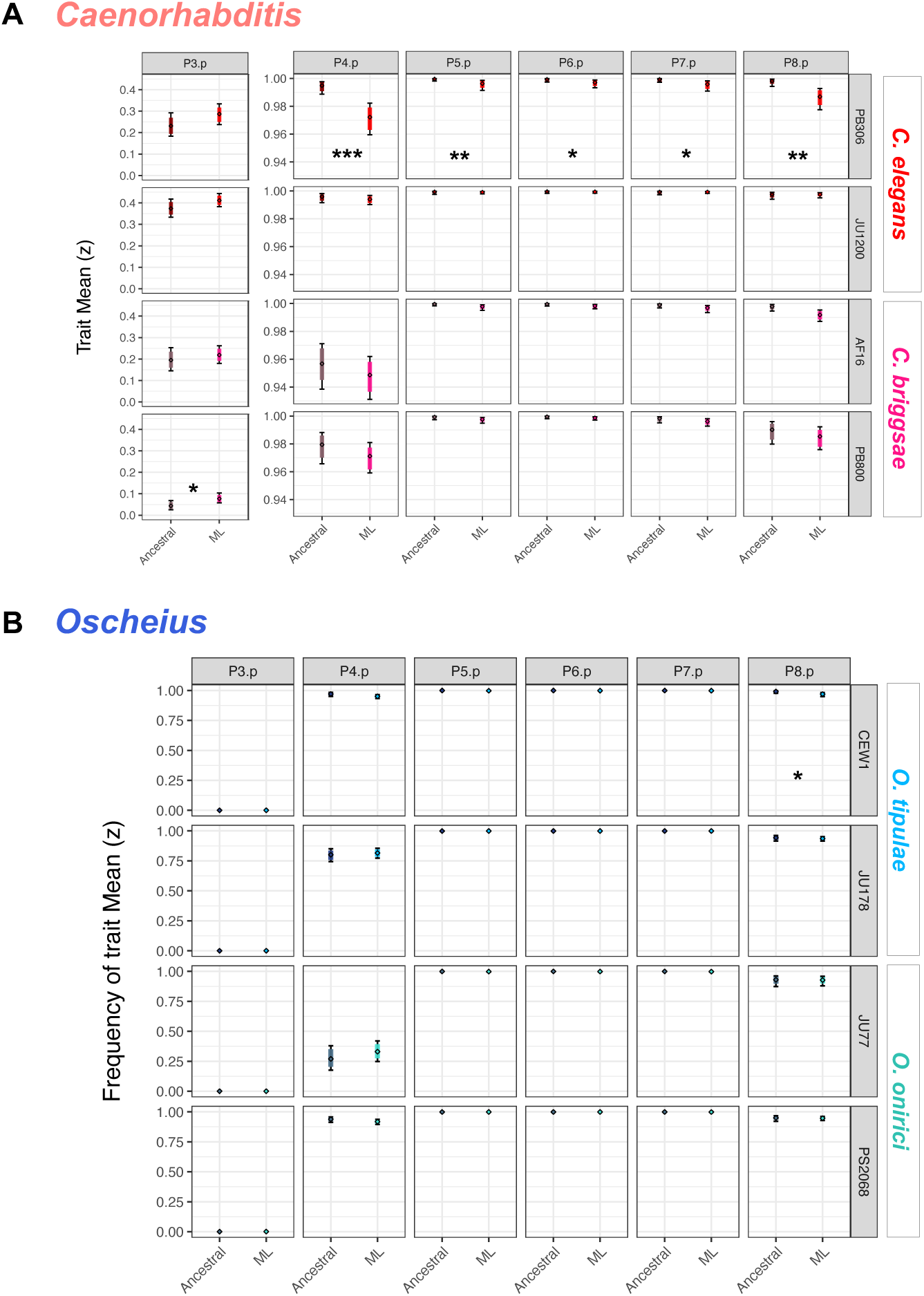
Mean mutational effects in *Caenorhabditis* and *Oscheius* species on vulval traits. Estimation of trait means (z) of VPC in ancestral pseudo-line (Ancestral, darker colours) and random mutant lines (ML, lighter colours) in two wild isolates of (**A**) *C. elegans* (PB306 and JU1200, red), *C. briggsae* (AF16 and PB800, pink), and (**B**) *O. tipulae* (CEW1 and JU178, blue) and *O. onirici* (JU77 and PS2068, turquoise). Each estimation displays the median (black circle), 83% (coloured bar) and 95% (black bar) credible intervals of the posterior distributions of the trait means in the observed data scale. Each posterior distribution has a minimum effective sample size of 2000. Significant MCMC *p*-values are denoted by * <0.05, ** <0.01, and *** <0.001. The scale of trait means of P3.p in *Caenorhabditis* species is adjusted between 0.0 and 0.4 division frequency.

**Figure S15.**
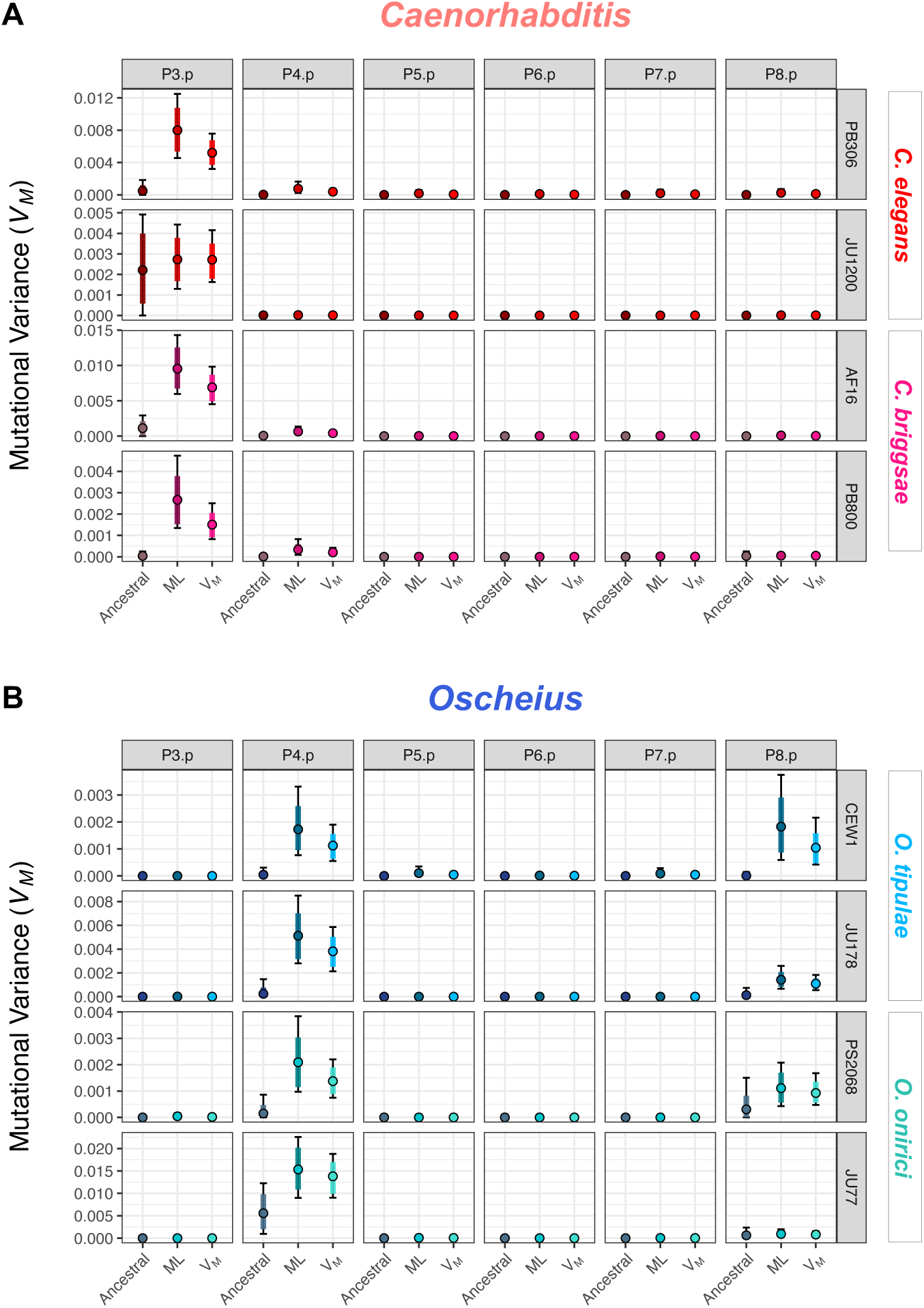
Mutational variance of vulval cells in isolates of *Caenorhabditis* and *Oscheius* species. Estimation of the mutational genetic variances (*V_M_*) of the six VPCs in two wild isolates of (**A**) *C. elegans* (PB306 and JU1200, red), *C. briggsae* (AF16 and PB800, pink), and (**B**) *O. tipulae* (CEW1 and JU178, blue) and *O. onirici* (JU77 and PS2068, turquoise). For each VPC are shown: left (Ancestral), variation among the ancestral pseudo-lines; middle (ML), variation among random mutant lines; right (*V_M_*), full *V_M_* model with a fixed effect named *Treatment* for control vs. mutant lines. The *V_M_* scale is adjusted to each isolate. Each estimation displays the median (black circle), 83% (coloured bar) and 95% (black bar) credible intervals of the posterior distributions of the genetic variances in the observed data scale. Each posterior distribution has a minimum effective sample size of 2000.

**Figure S16.**
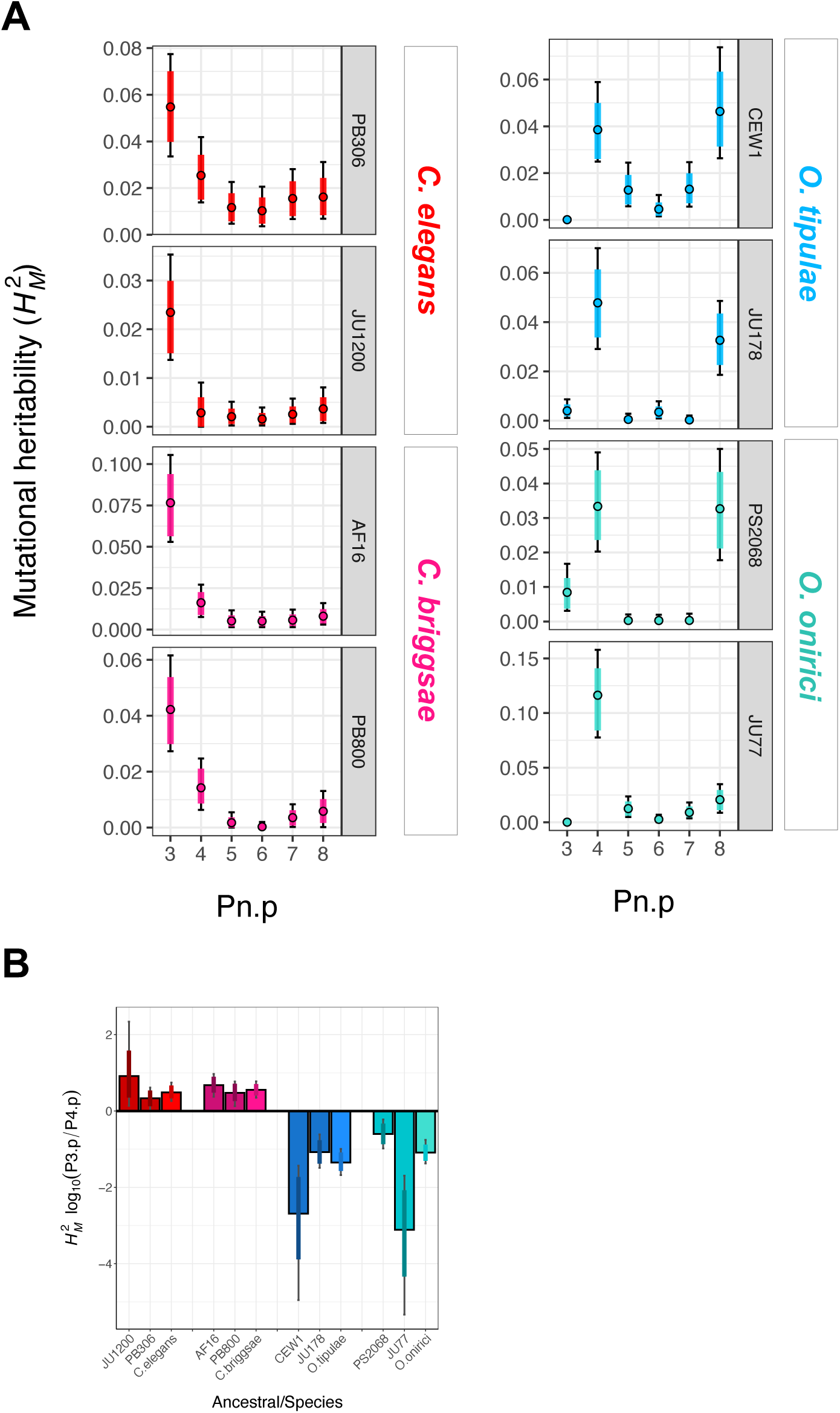
Mutational heritability of vulval cells in isolates and species of *Caenorhabditis* and *Oscheius*. Estimation of the broad-sense mutational heritability (*H*^2^*_M_*) of the six VPCs in (**A**) isolates of *C. elegans* (PB306 and JU1200, red), *C. briggsae* (AF16 and PB800, pink), *O. tipulae* (CEW1 and JU178, blue) and *O. onirici* (JU77 and PS2068, turquoise). The *H*^2^*_M_* scale is adjusted to each isolate and species. **(B)** Ten-fold change of the broad-sense mutational heritability (*H*^2^*_M_*) ratio between P3.p and P4.p in *C. elegans* (PB306 and JU1200, red), *C. briggsae* (AF16 and PB800, pink), *O. tipulae* (CEW1 and JU178, blue) and *O. onirici* (JU77 and PS2068, turquoise). Each estimation displays the median (bar plot), 83% (coloured bar) and 95% (black bar) credible intervals of the posterior distributions in the observed data scale with a minimum effective sample size of 2000.

**Figure S17.**
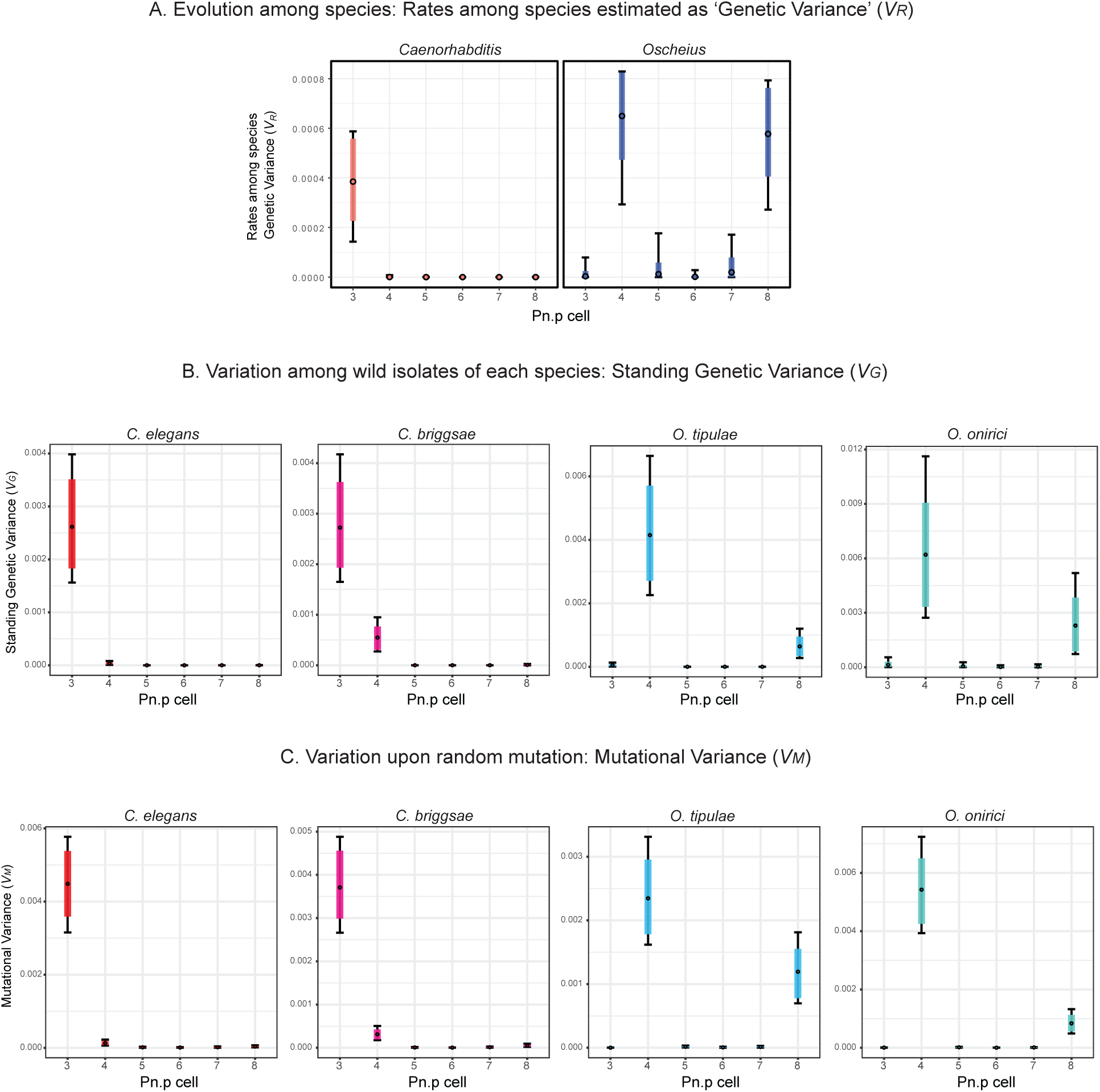
Contrasting vulva genetic variances in two nematode genera across scales. (A) Estimation of the rate among species genetic variance (*V_R_*) of the six VPCs in *Caenorhabditis* (salmon) and *Oscheius* (dark blue) genera. *V_R_* quantifies phenotypic evolutionary rates by estimating the increase of genetic variance among species per million years assuming a Brownian motion model of trait evolution. (B) Estimation of the standing genetic variances (*V_G_*) of the six VPCs in *C. elegans* (red), *C. briggsae* (pink), *O. tipulae* (blue) and *O. onirici* (turquoise) from the variation among wild isolates adjusted for the intrastrain variation of two APL controls on each species. (C) Estimation of the mutational variances (*V_M_*) of the six VPCs (P3.p to P8.p) in *C. elegans* (red), *C. briggsae* (pink), *O. tipulae* (blue) and *O. onirici* (turquoise). The mutational variance is the genetic variance among random mutation lines of two isolates corrected for their ancestral pseudo-line (APL) controls. The scale of the genetic variance (y-axis) is the same for both graphs in (A) and customised to each species in (B) and (C). Each estimation displays the median (black circle), 83% (coloured bar) and 95% (black bar) credible intervals of the posterior distributions of the genetic variances in the observed data scale. Each posterior distribution has a minimum effective sample size of 2000.

**Table S3.**

| Genus | Species | Phenotyped Nematodes |
| --- | --- | --- |
| *Caenorhabditis spp.* | 53 | 5347 [a] |
| *Oscheius spp.* | 24 | 2450 [b] |
| Total | 77 | 7797 |

Sample size of phenotyped *Caenorhabditis* and *Oscheius* species. Species were phenotyped in two replicates of 50 specimens each, except: [a] NIC394 (50+47) and NKZ352 (3Rx50); [b] JU75 (3Rx50). R: replicate.

**Table S6.**

| Species | Wild Isolates | Phenotyped Nematodes |
| --- | --- | --- |
| *C. elegans* | 50 | 5050 [a] |
| *C. briggsae* | 50 | 5000 |
| *O. tipulae* | 62 | 6250 [b] |
| *O. onirici* | 16 | 1600 |
| Total | 178 | 17900 |

Sample size of phenotyped *C. elegans*, *C. briggsae*, *O. tipulae* and *O. onirici* wild isolates ( $V_G$ ). Wild isolates were phenotyped in two replicates of 50 specimens each, except: [a] EG4349 (3Rx50), [b] JU1061(3Rx50). R: replicate.

**Table S7.**

| Species | wild isolates | Ancestral Pseudo-Lines (APL) | Phenotyped APL Nematodes | Random Mutation Lines (RML) | Phenotyped RML Nematodes |
| --- | --- | --- | --- | --- | --- |
| *C. elegans* | JU1200 | 24 | 2550 [a] | 50 | 6100 [b] |
|  | PB306 | 26 | 2600 | 49 | 4862 [c] |
| *C. briggsae* | AF16 | 24 | 2400 | 50 | 5000 |
|  | PB800 | 22 | 2200 | 50 | 5000 |
| *O. tipulae* | CEW1 | 18 | 1950 [d] | 50 | 5950 [e] |
|  | JU178 | 20 | 2000 | 50 | 5000 |
| *O. onirici* | PS2068 | 22 | 2200 | 50 | 5000 |
|  | JU77 | 16 | 1600 | 50 | 5000 |
| Total |  | 172 | 17500 | 399 | 41912 |

Sample size of phenotyped ancestral controls and mutation lines of *C. elegans*, *C. briggsae*, *O. tipulae* and *O. onirici* ( $V_M$ ). Strains were phenotyped in two replicates of 50 specimens each, except:
[a] JU1200 P\_PILOT (R1 150 + R2 100)
[b] JU3541/JU3554/ JU3557(R1 200 + R2 150); JU3608 (R1 200 + R2 150 + R3 50 + R4 50)
[c] JU3716 (R1 12 + R2 50)
[d] CEW1 P\_PILOT (R1 150 + R2 100)
[e] JU3499/ JU3501/ JU3525 (R1 200 + R2 150); JU3526 (R1 200 + R2 100)
R: replicate.

## Tables in separate files

**Table S1.** Nematode strains used in this work. The table contains several sheets: species, wild isolates, and mutation lines.

**(a)** List of *Caenorhabditis* and *Oscheius* species used to estimate rates of phenotypic evolution (*V_R_*).

**(b)** List of wild isolates of *C. elegans*, *C. briggsae*, *O. tipulae* and *O. onirici* used to estimate standing genetic variances (*V_G_*).

**(c)** List of Random Mutant Lines (RML)and respective ancestral in *C. elegans*, *C. briggsae*, *O. tipulae* and *O. onirici* used to estimate mutational variances (*V_M_*).

**(d)** *O. tipulae egl-20/Wnt* mutant.

**Table S2**. List and hyperlinks of nematode genome assemblies. The two sheets provide information including how to access the data.

**(a)** Rhabditina genome assemblies.

**(b)** Genome assemblies of *Caenorhabditis* and *Oscheius* species.

**Table S4.** Summary of the *V_R_* analysis in *Caenorhabditis* and *Oscheius* genera.

The rate among species genetic variance per million year (*V_R_*) was estimated among *Caenorhabiditis* and *Oscheius* species in the six vulva precursor cells (VPC), P3.p to P8.p. We estimated four different quantitative genetic parameters (Measure): Vr_Phylo (variance among-species dependent on the phylogeny), H2_Phylo (broad sense phylogenetic heritability), Vr_NotPhylo (variance among-species not dependent on the phylogeny), and trait_mean (trait mean). Each Measure is displayed in the liability (liab) and observed data (data) scales (Scale). For each posterior distribution is shown the mean, median, posterior mode, the 95% and 83% credible intervals, and effective size. All Pn.p cells were coded as binary trait, where the most common fate (Pnp_fate) for a given VPC was assigned a value of 1 and all others a value of 0. That is, in *Caenorhabditis* species the most common division fate for P3.p, P4.p and P8.p is one division (fate ‘SS’), while in *Oscheius* species it is two divisions (fate ‘SSSS’). For the central VPCs P5.p to P7.p, we considered their canonical 2°, 1°, 2° fates of each clade, respectively, as ‘wt’.

**Table S5.** Structure of phenotyping blocks to estimate vulval standing genetic variance (*V_G_*) and mutational variance (*V_M_*).

**Table S8.** Summary of the standing genetic variance analysis (*V_G_*). The two sheets provide the analysis for each genus.

**(a)** Summary of the posterior distributions of *V_G_* in *Caenorhabditis* species, and **(b)** in *Oscheius* species.

The standing genetic variance (*V_G_*) was estimated among wild isolates of *C. elegans*, *C. briggsae*, *O. tipulae* and *O. onirici* in the six vulva precursor cells (VPC), P3.p to P8.p. We estimated three different quantitative genetic parameters (Measure): vg (genetic variance among-lines), H2 (broad sense heritability), and trait_mean (trait mean). Each Measure is displayed in the liability (liab) and observed data (data) scales (Scale). Three different models (Model_set) are displayed: Control [among control Ancestral Pseudo-Lines (APL)], Wild (among wild isolates), and Vg (full model with APLs and wild isolates). For each posterior distribution is shown the mean, median, posterior mode, the 95% and 83% credible intervals, and effective size. All Pn.p cells were coded as binary trait, where the most common fate (Pnp_fate) for a given VPC was assigned a value of 1 and all others a value of 0. That is, in *Caenorhabditis* species the most common division fate for P3.p, P4.p and P8.p is one division (fate ‘SS’), while in *Oscheius* species it is two divisions (fate ‘SSSS’). For the central VPCs P5.p to P7.p, we considered their canonical 2°, 1°, 2° fates of each clade, respectively, as ‘wt’.

**Table S9.** Summary of the mutational variance analysis (*V_M_*). The four sheets provide the analysis for each genus, the first analysed for the four isolates of each genus, the second analysed per species.

**(a)** Summary of the Posterior distributions of *V_M_* in *Caenorhabditis* species with isolates independently, and **(b)** with the two isolates together for each species.

**(c)** Summary of the posterior distributions of *V_M_* in *Oscheius* species with isolates independently, and **(d)** with the two isolates together for each species

The mutational variance (*V_M_*) was estimated among Random Mutant Lines (RML) of *C. elegans*, *C. briggsae*, *O. tipulae* and *O. onirici* in the six vulva precursor cells (VPC), P3.p to P8.p. We estimated three different quantitative genetic parameters (Measure): m_vg (mutational genetic variance among-lines), H2 (broad sense mutational heritability), and trait_mean (trait mean). Each Measure is displayed in the liability (liab) and observed data (data) scales (Scale). Three different models (Model_set) are displayed: Control [among control Ancestral Pseudo-Lines (APL)], RML, and Vm (full model with APLs and RMLs). For each posterior distribution is shown the mean, median, posterior mode, the 95% and 83% credible intervals, and effective size. In all *V_M_* models, we ran models with isolates independently (a, c), and with the two isolates together for each species (b, d). All Pn.p cells were coded as binary trait, where the most common fate (Pnp_fate) for a given VPC was assigned a value of 1 and all others a value of 0. That is, in *Caenorhabditis* species the most common division fate for P3.p, P4.p and P8.p is one division (fate ‘SS’), while in *Oscheius* species it is two divisions (fate ‘SSSS’). For the central VPCs P5.p to P7.p, we considered their canonical 2°, 1°, 2° fates of each clade, respectively, as ‘wt’.

## Datasets in separate files

**Data S1**. Nematode vulva phenotyping species diversity.

Dataset of vulva phenotyping in *Caenorhabditis* and *Oscheius* species. We followed the vulva cell-fate terminology of (*84*). For nonvulval syncytial fates: S, no division; SS, one division; S ss, one division followed by a division of one of the daughters; ssss, two divisions. For vulval fates: U, undivided; t, transverse (left-right) division; l, longitudinal (antero-posterior) division; o, oblique division (between longitudinal and transverse); d, division (orientation could not be determined. Primes (‘) indicate the presence of an additional round of division. M is for missing cell.

**Data S2.** Phylogram of Rhabditina nematodes (nexus format).

**Data S3.** Phylogram of *Caenorhabditis* and *Oscheius* species (nexus format).

**Data S4**. Chronogram of *Caenorhabditis* and *Oscheius* species (nexus format).

**Data S5**. Chronogram of Rhabditina nematodes (nexus format).

**Data S6.** Vulva phenotyping for *V_G_* and *V_M_*.

Dataset of vulva phenotyping in *Caenorhabditis* and *Oscheius* species for control Ancestral Pseudo-Lines (CONTROL), Wild Isolates (WILD), and Random Mutant Lines. We followed the vulva cell-fate terminology of (*84*). For nonvulval syncytial fates: S, no division; SS, one division; S ss, one division followed by a division of one of the daughters; ssss, two divisions. For vulval fates: U, undivided; t, transverse (left-right) division; l, longitudinal (antero-posterior) division; o, oblique division (between longitudinal and transverse); d, division (orientation could not be determined. Primes (‘) indicate the presence of an additional round of division. M is for missing cell. In yellow are highlighted the deviations from the reference.

**Data S7.** Vulva phenotyping for *egl-20/Wnt* mutant in *O. tipulae*.

Dataset of parallel vulva phenotyping in *O. tipulae* wild-type (CEW1) and *egl-20/Wnt* mutant (JU152). We followed the vulva cell-fate terminology of (*84*). For nonvulval syncytial fates: S, no division; SS, one division; S ss, one division followed by a division of one of the daughters; ssss, two divisions. For vulval fates: U, undivided; t, transverse (left-right) division; l, longitudinal (antero-posterior) division. M is for missing cell. In yellow are highlighted the deviations from the reference.

